# Systems-level analysis of monocyte responses in inflammatory bowel disease identifies IL-10 and IL-1 cytokine networks that regulate IL-23

**DOI:** 10.1101/719492

**Authors:** Dominik Aschenbrenner, Maria Quaranta, Soumya Banerjee, Nicholas Ilott, Joanneke Jansen, Boyd A. Steere, Yin-Huai Chen, Stephen Ho, Karen Cox, Oxford IBD Cohort Investigators, Carolina V. Arancibia-Cárcamo, Mark Coles, Eamonn Gaffney, Simon Travis, Lee A. Denson, Subra Kugathasan, Jochen Schmitz, Fiona Powrie, Stephen Sansom, Holm H. Uhlig

**Author notes:** Corresponding author: Holm H. Uhlig, Translational Gastroenterology Unit, Experimental Medicine, University of Oxford, John Radcliffe Hospital Oxford, OX3 9DU, UK. These authors contributed equally to this work (Dominik Aschenbrenner, Maria Quaranta, Stephen Sansom, Holm H. Uhlig).

## Abstract

**BACKGROUND & AIMS:** Dysregulated immune responses are the cause of inflammatory bowel diseases. Studies in both mice and humans suggest a central role of IL-23 producing mononuclear phagocytes in disease pathogenesis. Mechanistic insights into the regulation of IL-23 are prerequisite for select IL-23 targeting therapies as part of personalized medicine.

**METHODS:** We performed transcriptomic analysis to investigate IL-23 expression in human mononuclear phagocytes and peripheral blood mononuclear cells. We investigated the regulation of IL-23 expression and used single-cell RNA-sequencing to derive a transcriptomic signature of hyper-inflammatory monocytes. Using gene network correlation analysis, we deconvolve this signature into components associated with homeostasis and inflammation in patient biopsy samples.

**RESULTS:** We characterized monocyte subsets of healthy individuals and patients with inflammatory bowel disease that express IL-23. We identified auto- and paracrine sensing of IL-1α/IL-1β and IL-10 as key cytokines that control IL-23-producing monocytes. Whereas Mendelian genetic defects in IL-10 receptor signalling induced IL-23 secretion, uptake of whole bacteria induced IL-23 production *via* acquired IL-10 signalling resistance. We found a transcriptional signature of IL-23-producing inflammatory monocytes that predicted both disease and resistance to anti-TNF therapy and differentiated that from an IL-23-associated lymphocyte differentiation signature that was present in homeostasis and in disease.

**CONCLUSION:** Our work identifies IL-10 and IL-1 as critical regulators of monocyte IL-23 production. We differentiate homeostatic IL-23 production from hyper-inflammation-associated IL-23 production in patients with severe ulcerating active Crohn’s disease and anti-TNF treatment non-responsiveness. Altogether, we identify subgroups of patients with inflammatory bowel disease that might benefit from IL-23p19 and/or IL-1α/IL-1β-targeting therapies upstream of IL-23.

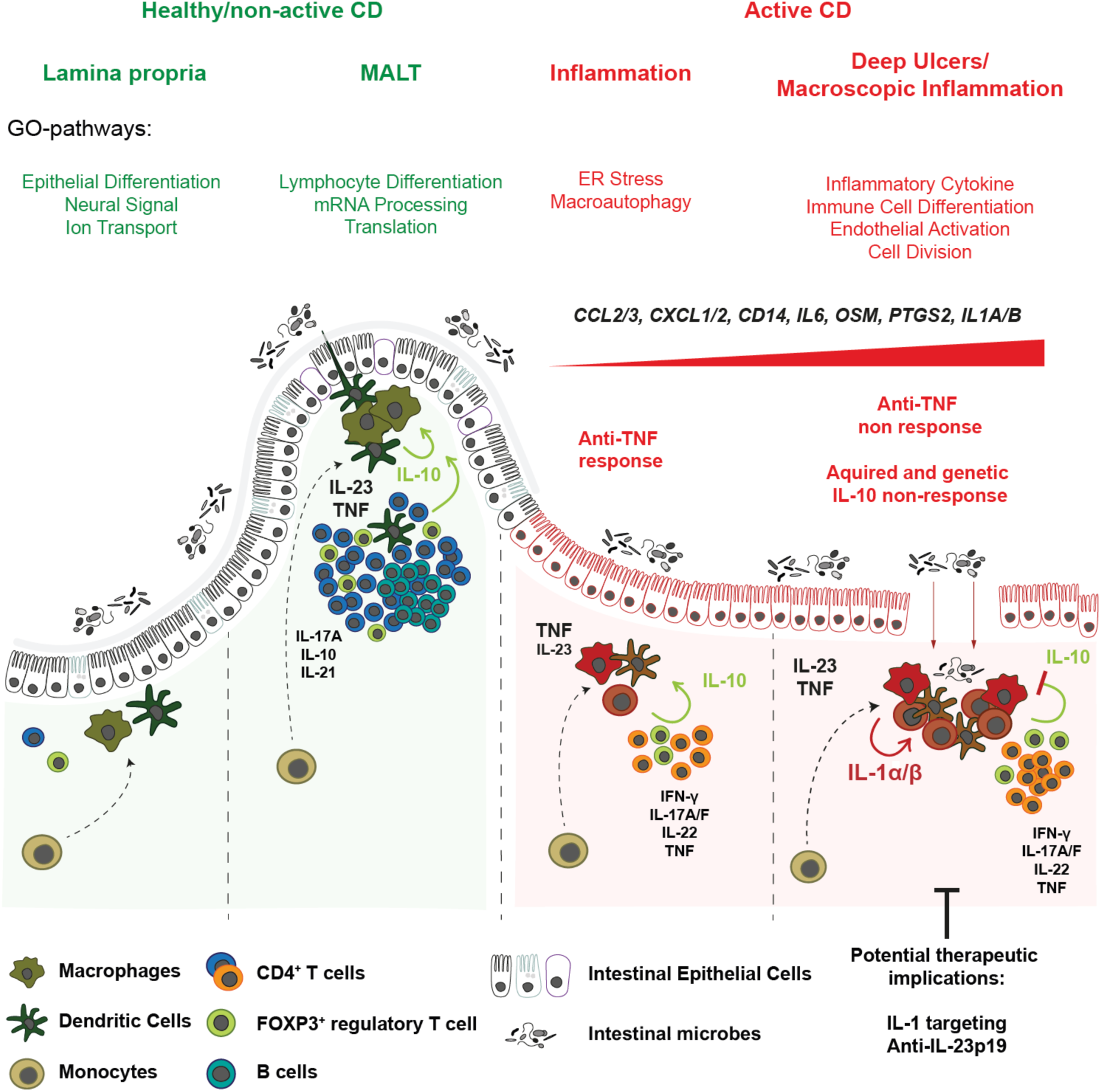

## INTRODUCTION

The pathogenesis of inflammatory bowel disease (IBD), that include Crohn’s disease (CD) and ulcerative colitis (UC) and IBD unclassified (IBDu) is caused by dysregulated innate and adaptive immune responses that drive chronic relapsing tissue inflammation^1^. Genetic studies suggest a complex polygenic inheritance driven by defective innate and adaptive immunity^1, 2^. IL-23 signalling and T-helper (Th)1/Th17 immunity are significant mediators of intestinal inflammation as indicated by genetic variants in *IL23R* encoding the IL-23 receptor^3^ as well as *RORC*, *STAT3*, *IRF5*, *IL1R1*, *IL6ST*, *IL12B, TYK2, IL21, JAK2, IFNG, SMAD3* and *CCR6*^4, 5^.

Monocytes, macrophages and dendritic cells produce IL-23 in response to bacteria- and fungi-derived microbial stimuli and drive Th1 and Th17 differentiation in the pathogenesis of intestinal inflammation^6–9^. In addition to a role of pro-inflammatory and anti-inflammatory cytokine networks^10^, mouse models as well as human Mendelian disorders highlight the essential role of IL-10 signalling in controlling inflammatory cytokine responses. Therapeutic approaches that target the pro-inflammatory arm of the immune system by blocking cytokine signalling or by affecting intestinal cell migration are effective in subgroups of patients with IBD^8^. Mice that lack IL-23p19 are protected from developing colitis in innate as well as lymphocyte replete models of intestinal inflammation induced by IL-10 signalling defects, bacterial colonisation or innate immune stimulation *via* the anti-CD40^7, 11^. Blockade of IL-23p19 and IL-12p40 showed therapeutic benefit for patients with CD and UC^12, 13^.

To predict genetic susceptibility or disease course, individual loci and genetic risk scores^14, 15^ as well as gene expression signatures or transcriptomic scores of peripheral CD8^+^ T cells^16^, peripheral blood^17^ or intestinal tissue^18^ are emerging patient stratification strategies in IBD. Membrane-bound TNF^19^ and IL-6/Oncostatin M (OSM) associated cytokines^18^ predict anti-TNF non-response. However, transcriptional signatures that implement IL-23 expression have not been described.

Personalised medicine targeting the IL-23 axis requires an understanding of the cellular sources, networks and regulation of IL-23. Here we investigate the regulation of IL-23, describe distinct monocyte subsets that express IL-23 and identify IL-1-signalling as the key cytokine for the differentiation IL-23-producing monocytes. We identify a hyper inflammatory signature of IL-23-producing monocytes in intestinal tissue transcriptomes of patients with IBD and find an additional signature of IL-23 that is associated with lymphocyte cell differentiation in healthy tissue.

## MATERIALS AND METHODS

### Human samples, cell isolation and cell culture

Patients with IBD and controls were recruited via the Oxford IBD cohort and Gastrointestinal biobank (REC 11/YH/0020 and 16/YH/0247). All patients with IBD and healthy volunteers provided written informed consent. Peripheral blood mononuclear cells (PBMC) were purified using Ficoll-Paque density gradient purification. For stimulation assays, 0.5×10^6^ -1×10^6^ PBMC or MACS-purified CD14^+^ monocytes (Mitenyi Biotec) were cultured in 200 μl medium in duplicates in round bottom 96-well plates. For detailed description of cell culture and stimulation, including whole bacteria preparation for stimulation, see Supplementary Experimental Procedures.

### Flow cytometry

We applied surface immunostaining for identification of immune cell subsets in complex mixtures of cells such as PBMC or enriched CD14^+^ monocytes. Surface staining and intracellular cytokine staining were combined for the analysis of frequencies of cytokine producing cells following stimulation and for the validation of single cell RNA-sequencing-identified monocyte clusters. Intracellular staining for phosphorylated STAT3 was used to analyse IL-10-responsiveness in CD14^+^ monocytes. For detailed description of experimental procedures (surface staining, intracellular cytokine staining, phosphoflow) stimulation conditions, reagents, antibodies and mathematical modelling of cytokine interactions please see Supplementary Experimental Procedures.

### Protein level analysis

Cytokine levels in cell culture supernatants were assayed using the Milliplex human cytokine/chemokine magnetic bead 41-plex panel (Milliplex Billerica, MA, USA; HCYTOMAG-60K-PX41) and acquired on a Luminex LX200 flow reader. For detailed description of experimental procedures please see Supplementary Experimental Procedures.

### ELISpot

A dualcolor ELISpot assay was developed for the detection of IL-10 and IL-23 producing cells. Please see Supplementary Experimental Procedures for detailed description of protocol and reagents.

### Single cell RNA-sequencing of monocytes, identification and characterisation of gene co-expression modules in the RISK cohort and Identification and characterisation of an IL-10-responsive monocyte gene signature

For a detailed description of cell isolation, stimulation, preparation of cells and single cell RNA-sequencing, data analysis including cross-condition analysis of single-cell RNA-sequencing data and comparison of LPS + anti-IL-10R single-cell RNA-sequencing data please see Supplementary Experimental Procedures.

### Patient cohorts and bioinformatics analysis

Cohorts of patients with IBD analysed in this study include the RISK study^20, 21^ GEO (GSE57945), Affymetrix microarray data from Janssen^22^ (GSE12251) and mucosal gene expression data by I. Arijs *et. al.*^23^ (GSE16879). For details regarding the analysis and data processing for the identification and characterisation of gene co-expression modules in the RISK cohort, identification and characterisation of an IL-10-responsive monocyte gene signature and validation see Supplementary Experimental Procedures.

### Gene expression analysis using real time PCR and expression array

Total RNA was extracted from cultured cells using the RNeasy Plus Mini Kit (Qiagen). The RNA yield was determined via Nanodrop ND1000 UV-vis Spectrophotometer. Complementary DNA (cDNA) was synthesized from 200 ng of total RNA and transcribed using the High Capacity cDNA Reverse Transcription Amplification Kit (Applied Biosystems). Real-time PCRs were performed in 96-well plates using the PrecisionPLUS qPCR Mater Mix (Primer Design) and the CFX96 Touch Real-Time PCR Detection System (BIO-RAD). The Affymetrix Human Transcriptome Array 2.0 (HTA2) was used for expression array analysis. For detailed description of methods, reagents and statistical analysis please see Supplementary Experimental Procedures.

### Statistical analysis

Statistical analyses were performed with GraphPad Prism, version 8.0 for Macintosh (GraphPad Software, La Jolla, CA) or Microsoft Excel for Mac, version 15.32. P-values ≤ 0.05 were considered significant and indicated as follows: *P ≤ 0.05; **P ≤ 0.01; ***P ≤ 0.001; ****P ≤ 0.0001. Statistical tests are described in figure legends.

## RESULTS

### IL-10 signalling blockade facilitates IL-23 production by a subset of peripheral blood monocytes

We investigated the regulation of *IL23A* expression by subjecting PBMC from 41 patients with IBD (**Supplementary Fig. 1A**; **Supplementary Table 1**) to IBD-relevant stimuli that target different aspects of innate and adaptive immune cell responses. Lipopolysaccharide (LPS), muramyl-dipeptide (MDP), T cell receptor and co-stimulation (αCD3/αCD28 coated beads) and IL-10 signalling blockade were used alone or in combination based on the concept that innate pathogen recognition receptor responses (in particular NOD2^14^ and TLR4^24^), T cell responses and IL-10 signalling defects are implicated by multiple genetic IBD susceptibility loci and Mendelian forms of IBD^25^. We performed microarray gene expression analysis of all conditions at 16 hours following stimulation in presence or absence of IL-10 receptor (IL-10R) blocking antibodies. We identified stimulus-specific and shared gene expression signatures (**Supplementary Fig. 1B and 1C**; **Supplementary Table 2**). LPS, L18MDP, and αCD3/αCD28 coated beads induced 319, 187 and 312 genes respectively out of which 109, 19, and 175 were condition-specific genes (BH adjusted p < 0.05, | fc | ≥ 1.5; **Supplementary Fig. 1B**). Changes in *IL23A* expression were moderate under these conditions (**Supplementary Fig. 1C**). We next investigated the role of IL-10 signalling during LPS, L18MDP, and αCD3/αCD28 stimulation. In total 36 genes were up- and 29 genes down-regulated by IL-10 blockade, with most of the changes being found during LPS stimulation (|fc| > 1.5, BH-adjusted p < 0.05; **Supplementary Fig. 1D and 1E**; **Supplementary Table 3**). *IL23A* expression was significantly up-regulated under conditions of LPS or L18MDP stimulation and IL-10 signalling blockade (**Supplementary Fig. 1C and 1D**).

**Figure 1:**
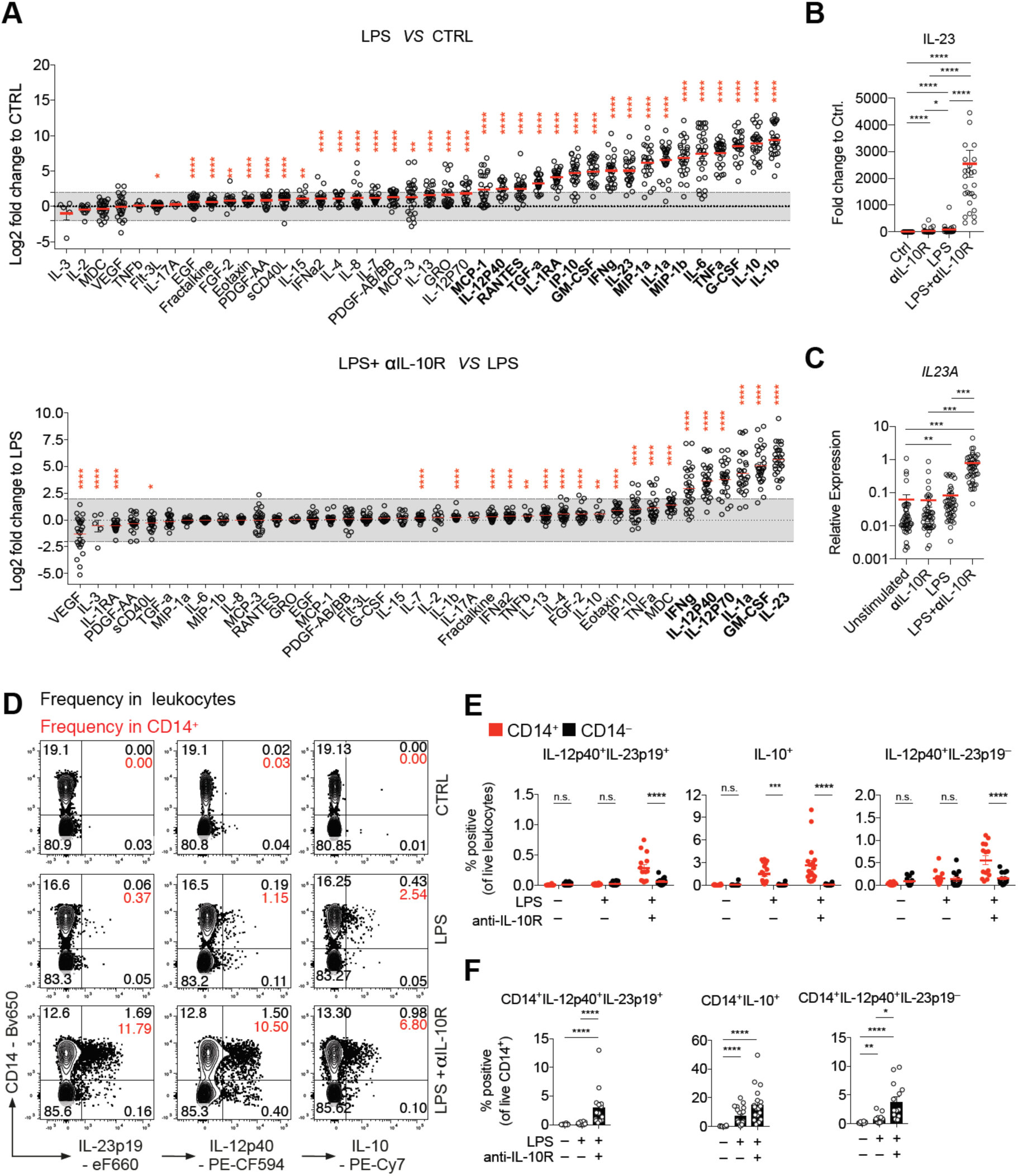
IL-10 regulates *IL23A* transcription and IL-23 protein secretion in a subset of monocytes from patients with IBD. (**A**) Analysis of PBMC culture supernatants collected after 16 hours stimulation with LPS +/– IL-10R blocking antibodies expressed as log2 fold change to unstimulated PBMC (CTRL) or LPS-stimulated PBMC (n = 28). Mean +/– SEM; Wilcoxon test, BH-adjusted p-values. (**B**) IL-23 protein concentrations in culture supernatants expressed as fold change to unstimulated PBMC (n = 28). Mean +/– SEM; Friedman test, BH-adjusted p-values. (**C**) RT-qPCR analysis of relative *IL23A* expression in PBMC (n = 45) following 16 hours stimulation. Mean +/– SEM; Kruskal-Wallis test, BH-adjusted p-values. (**D**) Contour plot presentation of IL-23p19-, IL-12p40- and IL-10-producing live leukocytes and CD14 surface expression measured at 16 hours post stimulation in PBMC. (**E**) Summary of frequencies of IL-12p40^+^IL-23p19^+^, IL-10^+^ and IL-12p40^+^IL-23p19^−^ CD14^+^ and CD14^−^ cells in total live leukocytes (n = 18). Mean +/– SEM; Mann-Whitney test. (**F**) Summary of frequencies of IL-12p40^+^IL-23p19^+^, IL-10^+^ and IL-12p40^+^IL-23p19^−^ of CD14^+^ cells (n = 18). Mean +/– SEM; Mann-Whitney test.

To investigate the regulation of secreted proteins by IL-10, we analysed cell culture supernatants after 16 hours stimulation (**Figure 1A**). LPS stimulation significantly upregulated protein secretion in 17 among the 40 proteins tested (| fc | ≥ 4-fold, BH Adjusted p < 0.05). The addition of IL-10 blockade significantly up-regulated 6 of the LPS-induced factors (IL-23p19, GM-CSF, IL-1α, IL-12p70, IL-12p40 and IFN-*γ*) (BH Adjusted p < 0.05, | fc | ≥ 4-fold) (**Figure 1A**). Compared to control, LPS stimulation resulted in a mean 85.23-fold induction (mean concentration control: 3.30 pg/ml, mean concentration LPS 135.98 pg/ml) of IL-23 (BH adjusted p < 0.05) while combined LPS and anti-IL-10R treatment induced a mean 2,544.95-fold increased (mean concentration LPS and anti-IL-10R 4737.57 pg/ml) IL-23 secretion (BH adjusted p < 0.05) (**Figure 1B**).

We next sought to understand the kinetics and cellular sources of IL-10 regulated LPS-responsive cytokines within the complex PBMC mixture. We quantitated IL-1α, IL-1β, IL-4, IL-6, IL-10, IL-12p40, IL-13, IL-17A, IL-23p19, GM-CSF, IFN-*γ* and TNF production in monocytes, NK cells, CD4^+^ T cells and CD8^+^ T cells by intracellular flow cytometry in patients with IBD and healthy donors (HD) PBMCs (**Supplementary Fig. 2A and 2B**). To capture the early and late phase of cytokine production we analysed cells at 6 and 16 hours following stimulation. At both time points of stimulation, CD14^+^ monocytes were the major source of IL-10, IL-1α, IL-1β, IL-6 and TNF (**Supplementary Fig. 2A and 2B**). CD4^+^ T cells contributed to the early cytokine response *via* expression of IFN-*γ*, IL-17A, GM-CSF and TNF, while CD8^+^ T cells produced IFN-*γ* and TNF (**Supplementary Fig. 2A and 2B**). NK cells contributed to the cytokine response by production of IFN-*γ* and TNF only at the late time point. At 16 hours stimulation monocytes still produced IL-1α and IL-1β while IL-6, TNF and IL-10 expression were reduced. All these cytokines were increased when cells were stimulated with LPS and anti-IL-10R. IL-23 (IL-12p40^+^IL-23p19^+^) and IL-12 (IL-12p40^+^IL-23p19^−^) expression were selectively increased in CD14^+^ relative to CD14^−^ leukocytes after IL-10 receptor blockade (**Figure 1D**, 1E **and 1F**). These results were confirmed in PBMC obtained from HDs (**Supplementary Fig. 2C, 2D and 2E)**.

**Figure 2:**
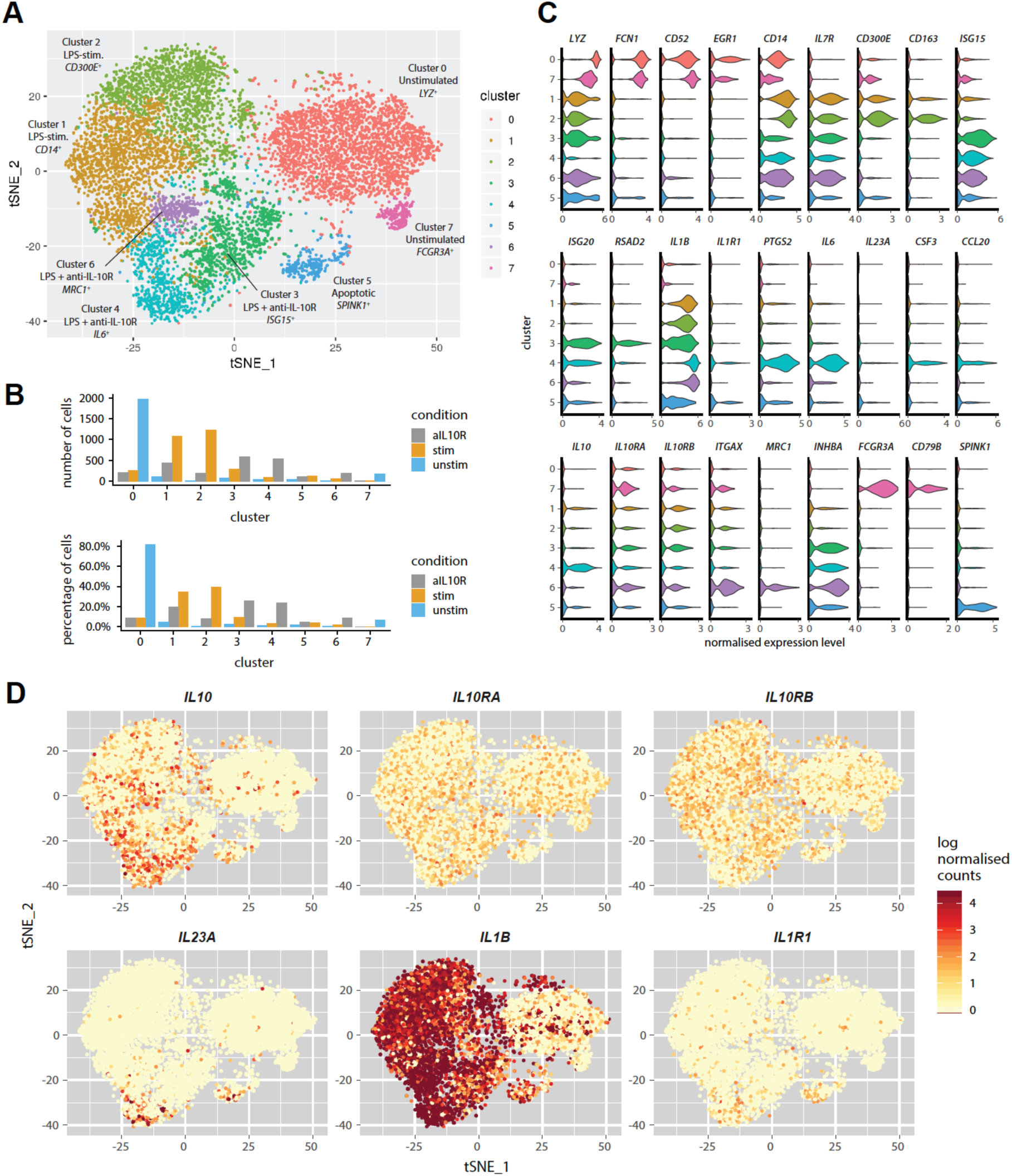
Single cell RNA-sequencing identifies subsets of inflammatory monocyte. (A) The tSNE plot shows the sub-populations of unstimulated, LPS-stimulated and combined LPS and anti-IL-10R stimulated monocytes that were identified by a graph-based clustering approach following cross-condition alignment with Harmony^50^. (**B**) The bar plots show the number and frequency of cells from the three stimulation conditions in each of the identified clusters. (**C**) The violin plots showing expression (x-axis) of genes characteristic of the identified monocyte clusters (y-axis). (**D**) Expression of *IL10, IL10RA, IL10RB, IL23A, IL1B and IL1R1* across the single monocytes according to figure (A).

Together, these results demonstrate that IL-10 signalling regulates IL-23 (IL-12p40^+^IL-23p19^+^) production in a subset of CD14^+^ monocytes (IBD mean = 3.03 % (95% CI of the mean: lower = 1.11, upper = 4.96); HD mean = 5.32 % (95% CI of the mean: lower = 3.51, upper = 7.14)).

### Single cell sequencing identifies inflammatory IL-10 regulated monocyte phenotypes

FACS analysis suggested the presence of unknown functional heterogeneity within the population of stimulated CD14^+^ monocytes (see *e.g.* **Figure 1D**). To characterise the LPS-induced and IL-10-regulated transcriptional profile of monocytes subsets, and to differentiate population-wide transcriptional changes from subset-specific responses we performed single cell RNA-sequencing (scRNA-seq) of unstimulated, LPS-stimulated and LPS and anti-IL-10R-treated CD14^+^ MACS-sorted monocytes from HDs (**Figure 2A**). Among the 8 clusters of monocytes that were detected, 2 clusters emerged after LPS stimulation and 3 additional clusters largely comprised of LPS and anti-IL-10R stimulated cells (**Figure 2A** **and** 2B).

The eight clusters showed discrete gene expression (**Figure 2C**) and were enriched for biological processes suggestive of different functional specialisations (**Supplementary Fig. 3A)**. The majority of unstimulated monocytes were lysozyme*^+^* (*LYZ*), *CD52^+^* (*CAMPATH1*), and *FCN1*^+^ expressing *CD14*^high^ monocytes (cluster 0 – unstimulated, classical monocytes), while a minority of unstimulated monocytes displayed a *CD52*^+^, *FCN1*^+^, *CD16*^+^ (*FCGR3A*) *CD14*^low^ phenotype (cluster 7 – unstimulated, non-classical monocytes). Two clusters of cells that emerged following LPS stimulation were comprised of *IL1B^+^* cells. The first showed expression of *CD14* and was enriched for genes associated with “T cell tolerance induction” (cluster 1 – LPS-stimulated), while the second displayed high *CD300E* and *CD163* expression and an enrichment for “monocyte activation” genes (cluster 2 – LPS-stimulated). The three additional monocyte phenotypes that appeared upon combined LPS stimulation and IL-10R blockade were demarcated by (i) high expression of type 1 interferon responsive genes (*e.g. IFIs*, *IFITs*, *ISGs*, *OASs, IRFs*) and genes linked to antigen processing and presentation (*e.g. B2M*, *CCR7*, *HLA, CD74*) (cluster 3 – LPS and anti-IL-10R, called IFN-induced monocytes); (ii) high expression of *ITGAX* and enrichment for genes associated with “superoxide generation” and “positive regulation of neutrophil activation” (cluster 6 – LPS and anti-IL-10R, called microbicidal monocytes) and (iii) expression of pro-inflammatory genes including (*IL1B*, *IL6*, *IL23A*, *CCL20* and *PTGS2*) and enrichment of genes associated with “Th17 cell lineage commitment”, “positive regulation of acute inflammatory response” and “interferon gamma production” (cluster 4 – LPS and anti-IL-10R; called IL-23^+^ inflammatory monocytes). These three phenotypes in LPS-stimulated anti-IL-10R-treated monocytes were replicated in a second donor (**Supplementary Fig. 3B**).

Interestingly, while *IL23A* and *IL1R1* were expressed in a cluster-specific manner under the “hyper-inflammatory” LPS and anti-IL-10R condition, *IL10, IL10RA* and *IL10RB* as well as *IL1A/IL1B* mRNA showed broader, cross-condition and cross-cluster expression (**Figure 2D**).

### IL-10-producing monocytes control IL-23-producing monocytes through paracrine signalling

In PBMC we observed that *IL23A* and *IL10* mRNA expression were strongly correlated (Spearmans’ r=0.84, p<0.0001) when stimulated with LPS and IL-10R blockade. **(Supplementary Fig 4A)**. These data indicate that *IL10* mRNA expression itself is tightly regulated by IL-10 signalling as part of a negative feedback mechanism. We assumed that *IL23A* and *IL10* would be produced by the same cells. However, inspection of the single-cell data suggested that this might not be the case and intracellular flow cytometry demonstrated that monocytes expressed IL-10 prior to up-regulating IL-23 (**Supplementary Fig. 2A and 2B and Supplementary Fig. 4B**) and that IL-10 and IL-23 largely originate from distinct cells (**Figure 3A**). This indicated that IL-10 may regulate IL-23 production through a paracrine mechanism in functionally distinct cells. Alternatively, *IL10* and *IL23A* mRNA expression might occur sequentially (*i.e.* early IL-10 producer become IL-23 producer) or an oscillatory fashion within individual cells.

**Figure 3:**
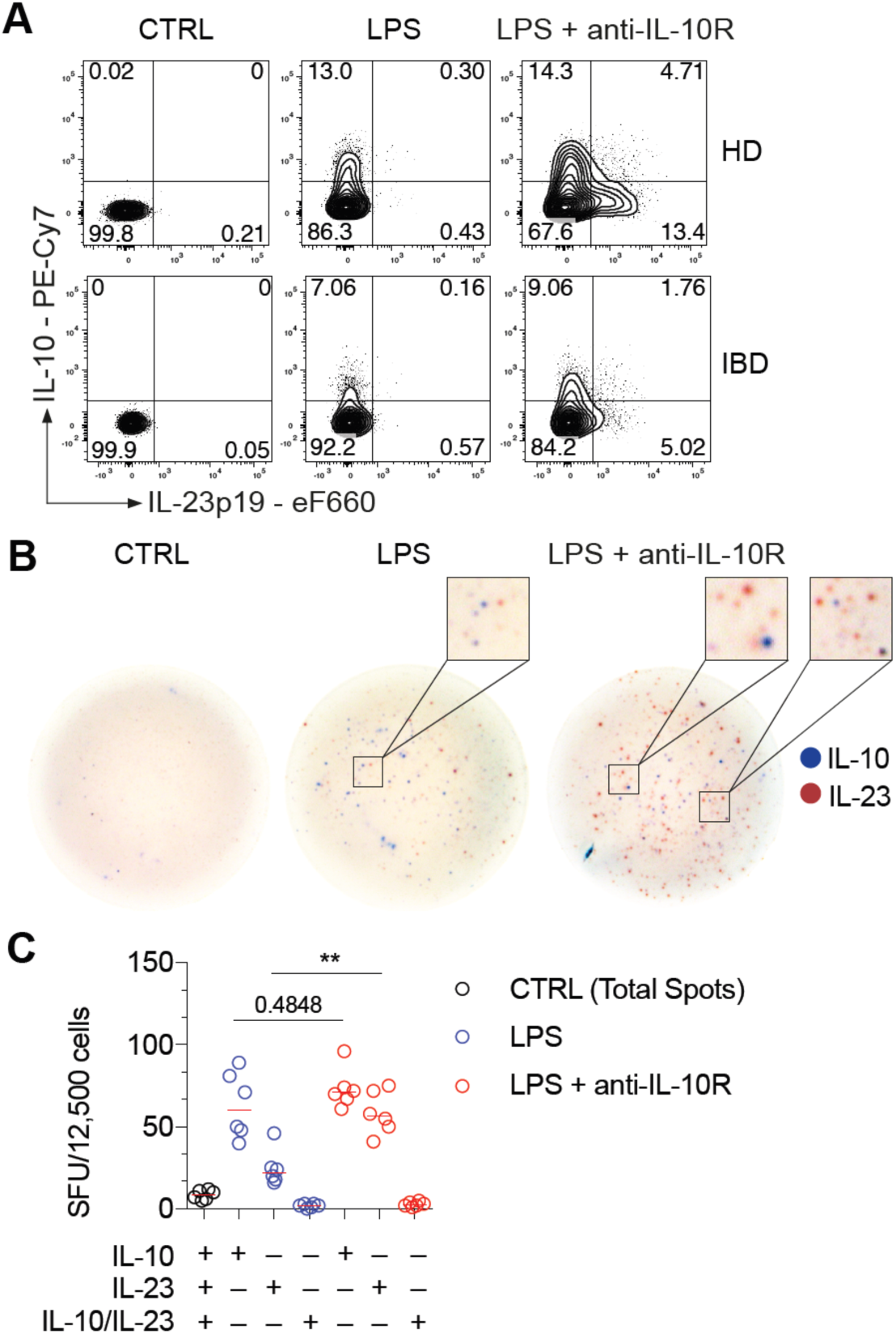
The monocyte response to PRR stimulation is heterogeneous revealing a paracrine mechanism of IL-10-depedent regulation of IL-23 production. (**A**) Representative contour plot presentation showing IL-23p19^+^ and IL-10^+^ frequencies in monocytes derived from a healthy donor and a patient with IBD at 16 hours post stimulation. (B) Representative dual-colour ELISpot images showing non-stimulated, LPS-stimulated and combined LPS and anti-IL-10R-stimulated monocytes IL-10 secretion (blue) and IL-12p40^+^IL-23p19^+^ secretion (red). (**C**) Summary of dual-colour ELISpot measurements from 3 independent experiments (n = 6, Mean +/– SEM; Mann-Whitney test).

**Figure 4:**
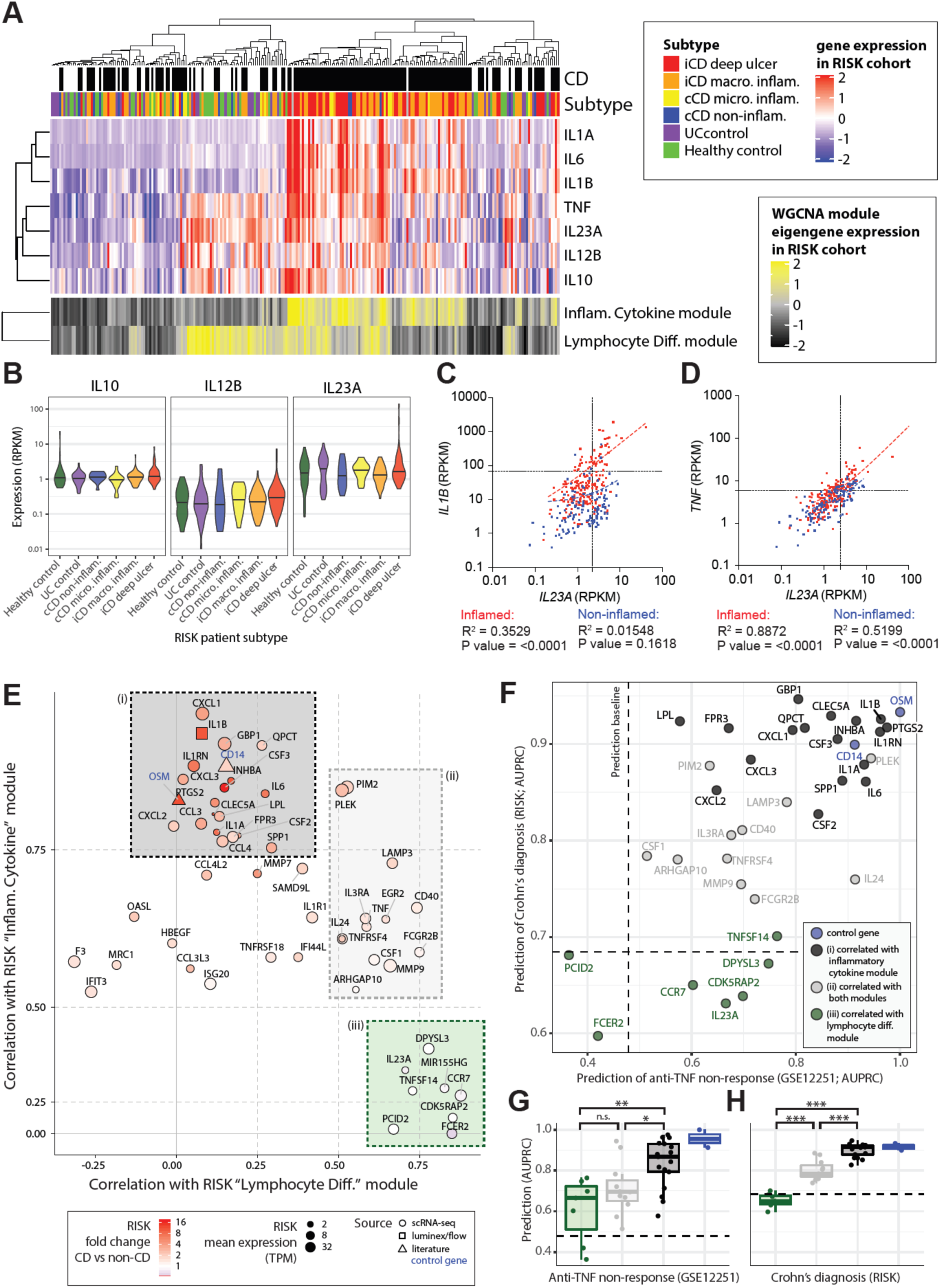
An IL-10-regulated inflammatory monocyte gene signature informs IL-23 and IL-1 targeting therapeutic approaches in inflammatory bowel disease. (**A**) Patients of the RISK cohort (diagnosis and subtype as shown in top panels) were clustered according to the expression of 22 modules of co-expressed genes (see Supplementary Figure 5A). The upper heatmap shows expression of key cytokines across the cohort. The lower panel shows expression of the eigen-genes of two of the identified modules of co-expressed genes. (**B**) The expression of *IL10*, *IL12B* and *IL23A* within the different patient strata of the RISK cohort. (**C** and **D**) Correlation of *IL23A* with *IL1B* (C) and *IL23A* with *TNF* (D) in the inflamed (red) and non-inflamed (blue) patient biopsies of the RISK cohort. (**E**) Correlation of genes specific to LPS and anti-IL-10R stimulated monocytes with the “inflammatory cytokine” and “lymphocyte differentiation” gene modules in the RISK cohort data identifies three subsets (shaded (i) black, (ii) grey and (iii) green boxes). (**F**) Assessment of the ability of the identified monocyte genes to predict anti-TNF non-response (x-axis, GSE12251) and diagnosis of Crohn’s (y-axis, RISK cohort). The dashed lines indicate random classifier performance. (**G** and **H**): comparison of the ability of the identified subsets of monocyte genes to predict *TNF* non-response in the Janssen cohort and diagnosis of CD in the RISK cohort (Wilcoxon tests, colours as shown in panels (E) and (F)). AUPRC: area under precision recall curve.

To distinguish between these possibilities, we performed a dual-colour ELISpot using MACS-purified monocytes. In line with a paracrine model where IL-10 producing monocytes regulate IL-23 production in others, we found that IL-10-producing monocytes were distinct from IL-23-producing monocytes and that double-stained cells were a minority (**Figure 3B** **and 3C**). These results suggest that IL-10 producing monocytes regulate IL-23 producing cells and demonstrate that the development of IL-23-producing monocytes is not a consequence of local cytokine segregation in tissue culture.

### Deconvolution of intestinal IL-23 gene expression in intestinal tissue

We next sought to understand the context of IL-23 expression during intestinal inflammation. We compared expression of IL-23 in ileal biopsies from CD patients and non-inflamed controls (HD or UC) in samples from the paediatric RISK cohort^20^ (**Figure 4A**). As expected, *IL23A* and *IL12B* were expressed in the CD patient samples along with *IL1A*, *IL1B* and *IL6*. However, we also found *IL23A* and *IL12B* expression in non-inflamed control biopsies (**Figure 4A** **and** 4B) where *IL23A* expression was associated with *TNF* rather than *IL1B* expression (**Figure 4C** **and** 4D). We therefore performed weighted gene co-expression network analysis (WGCNA)^26^ to investigate the possible sources and roles of *IL23A* in the inflamed and non-inflamed intestine (**Supplementary Fig. 5A**). This analysis identified 22 modules of co-expressed genes that were named according to their gene members and enrichments for biological pathways and sets of cell type marker genes (**Supplementary Table 4**). Clustering of the modules by their expression patterns in the RISK cohort patients identified six major groups that could be broadly distinguished by their correlations with diagnosis of CD, *IL23A* expression, epithelial gene expression, mitochondrial activity and cell growth genes. (**Supplementary Fig. 5A, Supplementary Fig. 5B**).

Most prominently, we found an “inflammatory cytokine” module that was significantly correlated with both CD (“crohns”, r=0.55) and *IL23A* expression (r=0.35) (**Supplementary Fig. 5B**). This module contained key myeloid and stromal markers genes (*CD14, PDPN*), pro-inflammatory cytokines (including *OSM, IL1B* and *IL6)* and Fibroblast activation protein (*FAP*) (**Supplementary Fig. 5B**) in keeping with the emerging concept of a pathogenic myeloid-stromal cell circuit in IBD^18, 24^. Intriguingly, however, we found that *IL23A* expression was most strongly correlated with two modules of “immune cell differentiation” (r=0.7) and “lymphocyte differentiation” (r=0.71) genes (**Supplementary Fig. 5A**). Of these, the “immune cell differentiation” module was weakly correlated with CD (r=0.2), associated with expression of myeloid (*CD14)* and lymphoid (*CD79A*, *CD4*) cell markers and enriched for “myeloid dendritic cell differentiation” (**Supplementary Fig. 5B and 6A**). The “lymphocyte differentiation” module showed an expression signature suggestive of lymphoid follicles including individual markers of B and T-cell identity (*CD79A*, *CD4*) and differentiation (*BACH2)*^25, 26^ (**Supplementary Fig. 5B**) as well as enrichments for gene ontology categories related to lymphoid cell differentiation, proliferation and selection (**Supplementary Fig. 6A**). While the “lymphocyte differentiation” module was not correlated with CD it was unique in showing significant enrichments for two sets of IBD GWAS associated genes (odds ratios > 3), suggesting that this set of genes is relevant for the pathogenesis of IBD (**Supplementary Fig. 6B**). We also noted that both of the *IL23A-*associated “immune cell differentiation” and “lymphocyte differentiation” modules were significantly enriched for genes belonging to the KEGG “Th17 cell differentiation” pathway (**Supplementary Fig 6A**).

**Figure 5:**
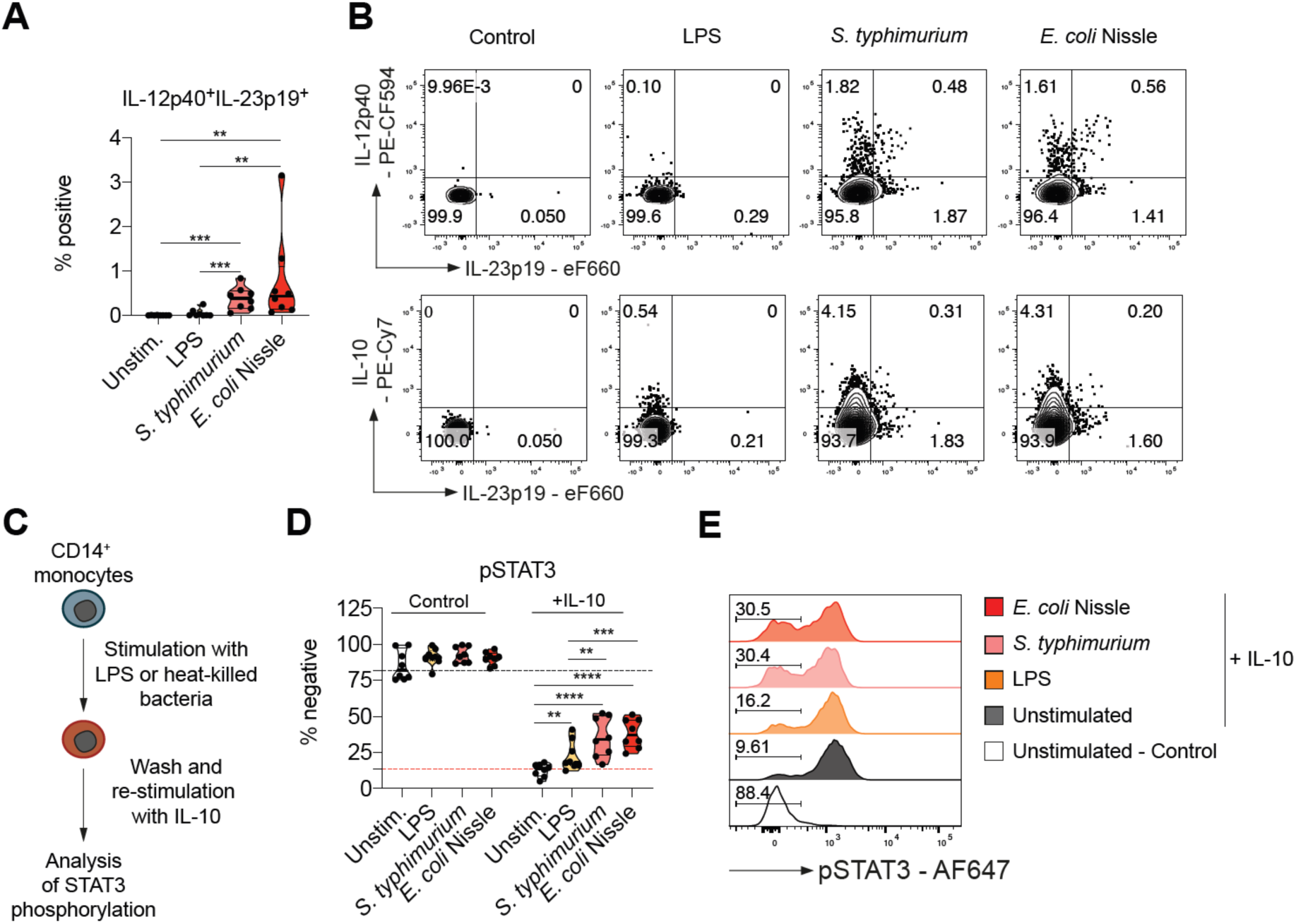
Monocyte uptake of whole bacteria causes IL-10 resistance and IL-23 secretion. (**A**) Frequencies of IL-12p40^+^IL-23p19^+^ live CD14^+^ at 16 hours following stimulation (n = 8, Mean +/– SEM; Mann-Whitney test). (**B**) Dot plots showing IL-12p40, IL-23p19 and IL-10 in live CD14^+^ monocytes from a healthy donor. (**C**) Scheme for the assessment of monocyte IL-10 responsiveness. (**D**) Frequencies of phospho-STAT3^−^ live monocytes without cytokine stimulation (Control) and following 15 minutes IL-10 (50 ng/µl) stimulation (n = 8, Mean +/– SEM; Mann-Whitney test). (**E**) Representative histograms showing phosphorylation of STAT3 in non-treated (Control) or IL-10-treated (+IL-10) live monocytes. Percentages of IL-10 stimulation-resistant monocytes are indicated.

**Figure 6:**
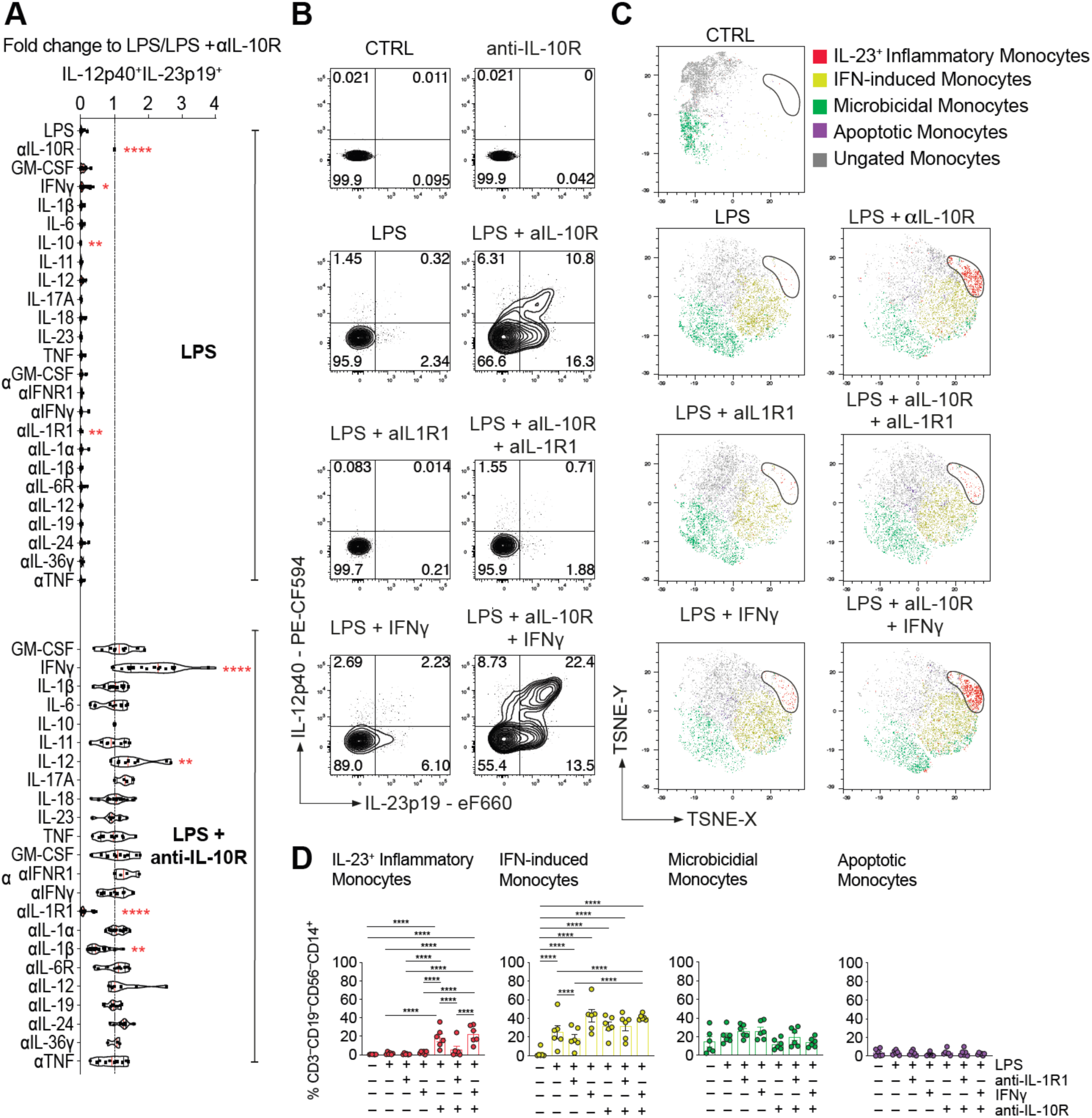
IL-1α and IL-1β signalling are essential for monocyte IL-23 production. PBMC from healthy donors (n = 4-20) were stimulated for 16 hours with combinations of LPS and αIL-10R in the presence of indicated exogenous human recombinant cytokine (all 10 ng/ml), and/or cytokine/cytokine receptor blockade (all antibodies 10 µg/ml). (**A**) Frequencies of IL-12p40^+^IL-23p19^+^ live CD14^+^ monocytes (Wilcoxon test; 95% CI). (**B**) Representative dot plot showing intracellular IL-12p40 and IL-23p19 according to (A). (**C**) tSNE presentation of IL-23p19, CCL20, HLA-DR, IDO-1, CCL2, S100A8, RPS6 and SPINK-1 expression in live CD14^+^CD3^−^CD19^−^CD56^−^–gated monocytes. Analysis of 3 healthy donors are shown as overlay. (**D**) Frequencies of monocyte clusters across stimulations based on cluster-specifying protein expression (n = 6; one way ANOVA after BH correction).

Overall, the network analysis suggested that *IL-23* expression was primarily associated with lymphoid cell differentiation in both health and disease. We therefore sought to understand which of the IL-10 regulated monocyte genes were specific to disease-associated inflammation. We correlated the expression of a curated set of IL-10 regulated monocyte genes (35 genes specific to combined LPS and anti-IL-10R stimulation scRNA-seq; **Figure 4E** **and** 4F; Supplementary methods section “Identification and characterisation of an IL-10-responsive monocyte gene signature”) with the eigengenes for the CD-associated “inflammatory cytokine” and the non-inflammatory “lymphocyte differentiation” modules in the RISK cohort data. This analysis identified three subsets of IL-10 regulated monocyte genes (**Figure 4E**). These comprised of (i) genes correlated only with the CD associated “inflammatory cytokine” module (black box, **Figure 4E**), (ii) a set of genes correlated with both the “inflammatory cytokine” and “lymphocyte differentiation” modules (grey box, **Figure 4E**), and (iii) genes correlated with only the non-inflammatory “lymphocyte differentiation” module (green box, **Figure 4E**). The IL-10-regulated genes that correlated with the “inflammatory cytokine” but not the “lymphocyte differentiation” eigengene (black box, **Figure 4E**) showed a significantly superior ability to predict anti-TNF response after 4 to 6 weeks treatment in an independent cohort of patients with UC (UC-cohort GSE12251)^22^ and diagnosis of CD in the RISK cohort^20^ (**Figure 4F**, 4G **and** 4H**, Supplementary Table 5**). By contrast, IL-10-regulated monocyte genes that correlated with “lymphocyte differentiation” but not “inflammatory cytokines”, had poorer predictive ability (**Figure 4F**, 4G **and** 4H). To validate our findings, we analysed intestinal transcriptome data of an additional adult cohort of patients with CD and UC (GSE16879)^22^. Similar to our initial results, inflammatory monocytes genes expression predicted disease activity and anti-TNF non-response in adult CD patients ileal biopsies transcriptomes. In colonic biopsies these genes were not predictive of disease activity but distinguished patients with CD or UC anti-TNF non-response (**Supplementary Fig. 7A and 7B**).

**Figure 7:**
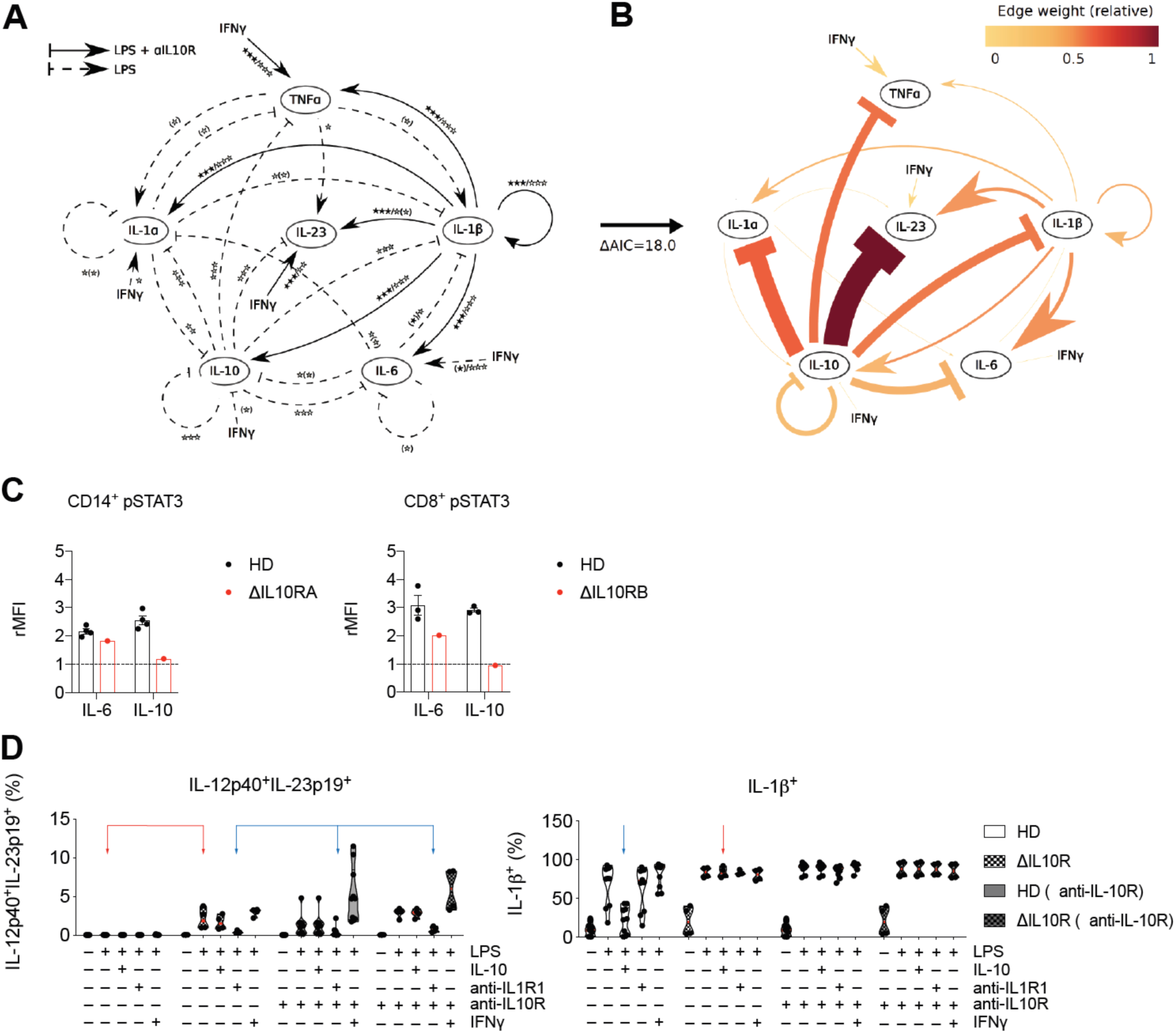
A mathematical model describing the modulation of monocyte IL-23 expression and analysis of monocyte IL-23 responses in patients with LOF in IL-10R1 and IL-10R2. (**A**) The extensive model represents the effects of the addition or blockade of cytokines in a PBMC culture in the presence of LPS or LPS and anti-IL-10R. Differences of cytokine addition or blockade in LPS stimulated samples (dashed arrows), in LPS and anti-IL-10R-stimulated samples (solid arrows), and in LPS and anti-IL-10R-stimulated and anti-IL-1β treated conditions (dotted arrows). Nominal effects with p>0.05 after false discovery rate (Benjamini & Hochberg) correction are shown in parenthesis. (**B**) A reduced complexity model was established by focusing on informative cytokine interactions. Edge weights are defined as the relative contribution to model fit and are dependent on the network configuration considered. The edges of the twenty-edge model have been coloured based on their weight. (**C**) Analysis of STAT3 phosphorylation following IL-6 or IL-10 stimulation in a patient with a loss of function variant in the gene coding for IL10RA (p.Y167C, left) and a patient with a loss of function variant in the gene coding for IL10RB (p.R117H, right). (**D**) Frequencies of IL-12p40^+^IL-23p19^+^ and IL-1β^+^ monocytes in PBMC from healthy donors and patients with IL-10R variants following 16 hours stimulation with combinations of LPS (200 ng/ml), IL-10 (10 ng/ml), anti-IL-10R (10 μg/ml), anti-IL-1R1(10 μg/ml) and IFN-γ (10 ng/ml).

These data therefore identify a subset of IL-10 regulated monocyte-derived genes, including *CXCL1/2*, *IL1A*, *IL1B*, *INHA*, *IL6*, *CCL3/4*, *PTGS2*, *CSF2/3* and *GBP1* that show a specific association for disease-associated intestinal inflammation in the RISK and GSE16879 CD cohort. By contrast, cytokines such as IL-23 and TNF were not specific for disease suggestive of context-dependent roles in homeostasis and inflammation.

### Monocyte uptake of whole bacteria causes IL-10 resistance and IL-23 secretion

The analysis of gene expression data in the RISK cohort suggests IL-23 expression despite presence of IL-10 in a subset of patients with macroscopic inflammation and deep ulcerating IBD in the absence of Mendelian IL-10 or IL-10R loss of function DNA variants, *i.e.* functional IL-10 resistance. In light of the epithelial barrier defect in ulcerating disease that predisposes to bacterial translocation, we tested whether direct contact of monocytes with bacteria can induce functional IL-10 resistance. We treated monocytes for 16 hours with-heat killed *Salmonella typhimurium* or *Escherichia coli* strain Nissle. Interestingly, monocytes produced IL-23 and IL-10 upon stimulation (**Figure 5A** **and** 5B) reminiscent of LPS stimulation in presence of IL-10R blockade. The cellular dichotomy of IL-23 and IL-10 production was maintained under these conditions (**Figure 5B**). To assess IL-10 responsiveness in previously stimulated monocytes we extensively washed cell cultures, subsequently exposed those to recombinant human IL-10 and evaluated phosphorylation of signal transducer and activator of transcription (STAT) 3 (**Figure 5C**). Indeed, bacterial stimulation was associated with an increased proportion of cells that did not respond to IL-10 demonstrating that monocyte encounter of whole bacteria induces a state of IL-10 resistance. (**Figure 5D** **and** 5E).

### IL-1α and IL-1β are essential for monocyte IL-23 production

The regulation of monocyte-derived IL-23 and IL-10 suggested co-regulation *via* additional factors (likely cytokines). We therefore investigated the functional effects of 11 co-regulated cytokines on monocyte IL-23 production (**Figure 6A** **and 6B, Supplementary Fig. 8A and 8B**). Strikingly, when blocking IL-1R1, IL-23 production was near completely abolished (**Figure 6A** **and** 6B**, Supplementary Fig. 8A-C**). Blocking IL-1β or IL-1α alone had only partial or no effect on IL-23 production indicating that either cytokine can compensate for the absence of the other in driving monocyte IL-23 production (**Figure 6A, Supplementary Fig. 8A**). The effect of IL-1β on IL-23 expression was context specific, since addition of IL-1β in the presence of anti-IL-10R blockade alone did not induce IL-23 expression **(Supplementary Fig. 8D).** None of the other tested cytokines (GM-CSF, IL-6, IL-11, IL-17A, IL-18, IL-19, IL-23, IL-24, IL-36*γ*, TNF and Type I Interferon) demonstrated a significant impact on IL-23 production. Although not essential, addition of IFN-*γ* or IL-12 increased monocyte IL-23 production (**Figure 6A** **and** 6B**, Supplementary Fig. 8A and 8B**).

To investigate the effects of IL-1R1 blockade on functional monocyte clusters, we stimulated PBMC from HDs with combinations of LPS and anti-IL-10R and analysed monocyte metaclusters-associated protein expression. In line with the single cell mRNA sequencing experiments the predicted clusters associated with LPS and combined LPS and anti-IL-10R stimulation (**Figure 6C, gating strategy outlined in Supplementary Fig. 8E)**. Blockade of IL-1R1 specifically inhibited the development of IL-23-producing monocytes while other monocyte clusters remained largely unaffected (**Figure 6C** **and** 6D). Importantly, monocyte IL-23 production induced by uptake of whole bacteria was similarly dependent on IL-1R1 signalling (**Supplementary Fig. 9A** and **9B**) suggesting that IL-23 is downstream of IL-1 signalling also in the context bacteria-induced functional IL-10 signalling defects.

### A mathematical model of IL-23 regulation in monocytes

To investigate the impact of cytokine networks on monocyte IL-23 production in more detail we analysed the effects of cytokines up-stream of IL-23 using a mathematical model of ordinary differential equations. Those describe interactions of monocyte-produced IL-23, TNF, IL-1α, IL-1β, IL-6, and IL-10 at the 16-hours time point. We identified an initial set of cytokine interactions, *i.e.* changes in cytokine production after addition or blockade of a cytokine (FDR corrected paired Wilcoxon test, **Figure 7A**). Then, we modelled all feasible network configurations, noting that most of the n=2^31^-1 configurations could be excluded a priori. We identified an optimal network describing the core network dynamics by ranking the models based on their fit to the data using the Akaike information criterion (**Figure 7B**). The model correctly identifies IL-10 signalling as negative feedback mechanism and IL-1β as a positive regulator (with IFN-*γ* as an amplifier) of IL-1α, IL-6 and IL-23 expression.

To confirm these predictions, we investigated the effect of IL-1R1 blockade on IL-23 expression in the context of genetic deficiency of IL-10 signalling. We stimulated PBMC obtained from patients with infantile onset IBD due to an *IL10RA* or *IL10RB* gene defect (**Figure 7C**). Interestingly, patient-derived monocytes produced IL-23 in response to LPS stimulation alone, indicating the intrinsic defect in IL-10R-dependent regulation, confirmed by the inability of exogenous IL-10 to suppress monocyte IL-1β production. Strikingly, IL-1R1 blockade inhibited monocyte IL-23 production in LPS-stimulated PBMC from IL-10R deficient patients (**Figure 7D**). Together, these analyses confirmed IL-10 as the major negative regulator of IL-23 production, while IL-1 signalling (and in particular IL-1β) is essential for monocyte IL-23 synthesis.

## DISCUSSION

We identify key regulatory circuits of IL-23 production by inflammatory monocytes. These include failure of paracrine IL-10-mediated control as well as autocrine and paracrine signalling of IL-1α and IL-1β in response to inflammatory stimuli. Indeed, increased IL-23 production can be observed in monocytes from patients with infantile onset Mendelian IL-10 signalling deficiency or experimental blockade of the IL-10 receptor, but also monocytes that respond to whole bacteria express IL-23 as a consequence of acquired IL-10 non-responsiveness. Most importantly, we define a transcriptional signature of IL-23^+^ inflammatory monocytes that indicates a state of acquired IL-10 non-responsiveness in a subgroup of patients with deep ulcers and compromised epithelial function where monocytes may directly respond to whole bacteria. This signature predicts both diagnosis of CD and resistance to anti-TNF treatment similar to previously described OSM expression^18^. Interestingly, loss of IL-10 responsiveness induces *IL23A* mRNA and IL-23 protein expression only in a small fraction of monocytes that express the IL-1 receptor *IL1R1*. This selective IL-23 expression contrasted with pervasive induction of IL-1α and IL-1β expression as well as a broad induction of IL-10 expression in monocytes under the same hyper-inflammatory condition.

Our data support context-specific roles of IL-23 in driving the differentiation of pathogenic Th17/Th1 cells in presence of IL-1^27^ that are enriched in inflamed intestinal tissue from patients with IBD^28, 29^, while supporting the development of non-pathogenic Th17 cells under homeostatic conditions such as those found in mucosa-associated lymphoid tissue (MALT) - in healthy individuals.

Together, these results suggest that only subgroups of patients with active disease may benefit from anti-IL-23p19 therapy, and that the targeting of upstream cytokines that regulate IL-23 expression in inflammation, such as IL-1 might provide a selective means to block pathogenic IL-23 expression.

The critical element that differentiates homeostatic IL-23 and TNF expression from hyper-inflammation is the co-expression of a multitude of IL-10-regulated factors, including IL-1α and IL-1β in patients with severe active disease and anti-TNF treatment non-responsiveness.

The strong predictive ability of an IL-10 sensitive inflammatory monocyte signature (*CD14*, *IL1A*, *IL1B, OSM*, *PTGS2*, *IL6*, *CCL2/3* and *CXCL1/2*) emphasises a surprising and underestimated extent of IL-10 non-responsiveness in IBD. In those patients inflammation is present despite transcription of IL-10 (patients express large amount of IL-10) and absence of pathogenic variants within the IL-10 receptor (exome sequencing did not reveal Mendelian forms of IL-10 receptor signalling defects in this cohort^30^). The high expression of *PTGS2* in this subgroup of patients with IBD represents an additional indicator for deregulated IL-10 responses and intestinal antimicrobial immunity^31^. Our experiments suggest that direct contact of monocytes with whole bacteria can induce IL-10 resistance that allows co-expression of IL-23 and IL-1 explain the signature of IL-10 non-responsiveness in a subgroup of patients with intestinal ulceration. An additional mechanism of reduced IL-10 responsiveness may be differential expression of IL-10R1^32^. Our ELISpot experiments exclude differential spatial clustering of IL-10 or IL-23 secreting monocytes as well as oscillatory or sequential IL-10 and IL-23 transcription. This suggests that the inflammatory microenvironment drives in parallel several monocyte effector programs and monocyte to macrophage and DC differentiation processes, reminiscent to plasmacytoid dendritic cells (pDC) stimulation with a single stimulus (R848 or CpG) of that induces diverse transcriptional states and cellular functions^33^. This is likely a biological mechanism to ensure functional heterogeneity under hyper-inflammatory conditions, when diversity in the host defence towards a range of potential pathogens is essential.

Single cell RNA transcriptomic approaches identified transcriptional signatures in circulating and tissue human monocytes and macrophages^9, 34–38^ demonstrating phenotypic and functional diversity in monocyte, macrophage and DC populations. We have focused on the cellular heterogeneity and inflammatory response of peripheral monocytes, because those cells are recruited into the gut and differentiate into pro-inflammatory MHC-II-high monocytes and macrophages that outnumber resident, yolk-sac derived tissue macrophages during inflammation^39–41^. Such pro-inflammatory macrophages have been characterized following *in vitro* differentiation using LPS-stimulation and express IL-12 and IL-23 in response to STAT1-dependent IFN-γ signalling^42^.

Our data suggest that the source of IL-23 in the context of inflammasome activation and IL-1 production in the inflamed intestine are inflammatory monocytes and CX3CR1^+^IL-1β^+^ positive macrophages^43, 44^. In individuals without intestinal inflammation, our network analysis suggested that *IL23A* expression was likely derived from terminal ileal MALT because the “lymphocyte differentiation” to which it was assigned also showed strong correlations with orthologs of genes associated with murine small intestine lymphoid tissue (SILT) including *IL22RA2*, *ITGAX*^27^ as well as with *VCAM1*, which is a marker of lymphoid associated villi in humans (**Supplementary Fig. 5B**). In such MALT tissue, which is hyperplastic in paediatric patients^45^ the source of *IL23A* is most likely dendritic cells (DC), tolerogenic CD103^+^DC^46^ or macrophages as is known to be the case in mouse SILT^43^. This work demonstrates the utility of network-based approaches for deconvolving variance in gene expression that arises from the analysis of biopsy samples from tissues with complex micro-anatomical heterogeneity such as the small intestine.

Our mathematical model pin-points IL-1 as a key positive regulator of IL-23 expression in monocytes. Studies in monogenic forms of IBD suggest that IL-1R1 blockade can resolve intestinal inflammation in patients with IL-10 receptor defects^47^, mevalonate kinase defects and potentially gain of function NLRC4^48^ defects, which are all characterised by increased inflammasome activation and IL-1 secretion. It is therefore an attractive hypothesis that the hyperinflammatory IL1^+^PTGS^+^IL6^+^ signature might define an anti-IL-1R1 responsive group of patients with IBD. Interventional studies are required to identify differential effects between IL-23 and IL-1 blockade since both have complex effects on licensing for effector cytokine production and differentiation of Th1 and Th17 cells^49^. Altogether our findings may inform transcriptional diagnostic efforts to guide use of IL-23p19 targeting therapies and suggest a role for therapeutics capable of selectively blocking IL-23 in disease by targeting context-specific upstream factors such as IL-1.

## Conflict of Interest Statement

This research was funded by the National Institute for Health Research (NIHR) Oxford Biomedical Research Centre (BRC). The views expressed are those of the author(s) and not necessarily those of the NHS, the NIHR or the Department of Health. The study has been supported via a collaborative grant by Eli Lilly. BAS, SH, KC and JS are current or previous employees of Eli Lilly. HHU received research support or consultancy fees from UCB Pharma, Eli Lilly, Boehringer Ingelheim, Pfizer and AbVie. FP has received research support or consultancy fees from GSK, UCB Pharma, Medimmune, Janssen and Eli Lilly. SPLT has been adviser to, in receipt of educational or research grants from, or invited lecturer for AbbVie; Amgen; Asahi; Biogen; Boehringer Ingelheim; BMS; Cosmo; Elan; Enterome; Ferring; FPRT Bio; Genentech/Roche; Genzyme; Glenmark; GW Pharmaceuticals; Janssen; Johnson & Johnson; Eli Lilly; Merck; Novartis; Novo Nordisk; Ocera; Pfizer; Shire; Santarus; SigmoidPharma; Synthon; Takeda; Tillotts; Topivert; Trino Therapeutics with Wellcome Trust; UCB Pharma; Vertex; VHsquared; Vifor; Warner Chilcott and Zeria. SK has received consultancy fees from Janssen and Takeda.

## Author contributions

DA, MQ performed experiments. SNS, SB, DA, NI, BAS, and JJ performed bioinformatics analyses. SNS was responsible for the scRNA-seq, WGCNA and signature deconvolution analyses. JJ, MC and EG were responsible for the mathematical modelling. CVA-C, ST, HHU, TD and SK contributed patient cohort recruitment and analysis. JS, ST, SK, FP, SNS and HHU supervised the study. All authors discussed data and contributed to the manuscript.

## Data availability

The data supporting the findings described in this study are available from the corresponding author upon request. Single cell RNA sequencing data generated in this study have been deposited at NCBI’s Gene Expression Omnibus (GEO) and are accessible under GEO Series accession number GSE130070. Microarray data generated in this study have been deposited at NCBI’s Gene Expression Omnibus and are accessible under GEO Series accession number GSE137680. Both data sets are available *via* a GEO SuperSeries that represents the publication as a whole (GSE138009).

## Abbreviations

BH: Benjamini & Hochberg
CD: Crohn’s disease
CD: Cluster of differentiation
FACS: Fluorescence assisted cell sorting
GFP: Green fluorescent protein
IBD: Inflammatory bowel disease
IBDu: IBD unclassified
IL: Interleukin
LPS: Lipopolysaccharide
MACS: Magnet-assisted cell sorting
MDP: Muramyl-dipeptide
OSM: Oncostatin M
PBMC: Peripheral blood mononuclear cells
scRNA-seq: single cell RNA-sequencing
STAT: Signal transducer and activator of transcription
Th: Thelper
TNF: Tumour necrosis factor
UC: Ulcerative colitis.

## Acknowledgments

We thank all patients, volunteers and blood donors for participation in this study. We thank Priya Siddhanathi, James Chivenga, Jennifer Hollis and Sebastian Rogatti Granados for excellent technical assistance, Helen Ferry for flow cytometry support. We thank the Oxford Genomics Centre Single-cell core unit for their technical assistance and support. We like to thank Jon Sedgwick for discussion. We acknowledge the contribution of the BRC Gastrointestinal biobank (11/YH/0020, 16/YH/0247), which is supported by the National Institute for Health Research (NIHR) Oxford Biomedical Research Centre (BRC). We acknowledge support of the BRC (FP, ST, HHU), the Leona M. and Harry B. Helmsley Charitable Trust (FP, HHU), research grants from the Crohn’s & Colitis Foundation of America (LAD, SK, HHU, FP and SP), the COLORS in IBD project via a ESPGHAN network grant (HHU), the Wellcome Trust (FP), the Kennedy Trust for Rheumatology Research (SNS), and to the individual study institutions participating in the RISK study. We thank the Oxford IBD Cohort Investigators** Adam Bailey, Ellie Barnes, Elizabeth Bird-Lieberman, Oliver Brain, Barbara Braden, Jane Collier, James East, Alessandra Geremia, Lucy Howarth, Satish Keshav, Paul Klenerman, Simon Leedham, Rebecca Palmer, Astor Rodrigues, Alison Simmons, Peter Sullivan. The views expressed are those of the author(s) and not necessarily those of the NHS, the NIHR or the Department of Health.

## SUPPLEMENTARY EXPERIMENTAL PROCEDURES

### Patient cohort - RISK cohort and outcome classification

The RISK study is an observational prospective cohort study with the aim to identify risk factors that predict complicated course in pediatric patients with Crohn’s disease^20^. The RISK study recruited treatment-naive patients with a suspected diagnosis of Crohn’s disease. The Paris modification of the Montreal classification were used to classify patients according to disease behaviour (non-complicated B1 disease (non-stricturing, non-penetrating disease); complicated disease, composed of B2 (stricturing) and/or B3 (penetrating) behaviour) as well as disease location (L1, ileal only, L2, colonic only, L3, ileocolonic and L4, upper gastrointestinal tract). 322 samples were investigated with ileal RNA-seq. Individuals without ileal inflammation were classified as non-IBD controls. Patients with Crohn’s disease were followed over a period of 3 years. Patients were largely of European (85.7%) and African (4.1%) ancestry.

### Cell culture and stimulation

For stimulation assays, 0.5×10^6^ -1×10^6^ PBMC or MACS-purified CD14^+^ monocytes (Mitenyi Biotec) were cultured in 200 μl medium in duplicates in round bottom 96-well plates and exposed to ultrapure 200 ng/ml LPS (Enzo Life Sciences; Cat.# ALX-581-008), 200 ng/ml L18MDP (Invivogen) 10 μg/ml anti-IL-10R (Biolegend; clone: 3F9), anti-CD3/anti-CD28 beads (Invitrogen) or Staphylococcal enterotoxin B (SEB; Sigma) for the indicated time in complete RPMI with L-glutamine (Sigma) supplemented with 10 % human serum (Sigma; Cat.# H4522), non-essential amino acids (Gibco); 1 mM Sodium-Pyruvate (Gibco) and 100 U/ml penicillin and 10 μg/ml streptomycin (Sigma). Supernatants were collected and stored at -80 °C for the quantification of cytokine production. The following neutralizing antibodies or receptor blocking antibodies were used (all 10 μg/ml): anti-IL-24 (R&D; Cat.# AF1965; Polyclonal Goat IgG), anti-IL-12p70 (R&D; Cat.# MAB219; clone 24910), anti-IL-19 (R&D; Cat.# MAB10351; clone 152107), anti-GM-CSF (R&D; Cat.# MAB215; clone 3209), anti-IL-1R1 (R&D; Cat.# AF269; polyclonal Goat IgG), anti-IL-1α/IL-1F1 (R&D; Cat.# AF-200-NA; polyclonal Goat IgG), anti-IL-1β/IL-1F2 (R&D; Cat.# MAB201; clone: 8516), anti-IL-6R (Tocilizumab, Actemra*®*, Roche,), anti-TNFα (Infliximab, REMICADE*®*, Janssen), anti-IFNα/βR2 (Millipore; Cat.# MAB1155; clone: MMHAR-2), anti-IL-10R (Biolegend; Cat.# 308807; clone: 3F9), anti-IL-10 (R&D; Cat.# MAB217; clone: 23738), anti-IFN*γ* (Biolegend; Cat.# 506513; clone: B27), anti-IL-36*γ* (R&D; Cat.# MAB2320; clone: 278706). The following recombinant human cytokines were used (all 10 ng/ml): IL-1β (Preprotech), IL-6 (Preprotech), IL-10 (Preprotech), IL-11 (R&D), IL-12 (Preprotech), IL-17A (Preprotech), IL-18 (R&D), IL-23 (Preprotech), GM-CSF (Preprotech), IFN*γ* (Preprotech), TNFα (Preprotech).

### Whole bacteria preparation and stimulation

Both *Salmonella enterica serovar Typhimurium* (ST) expressing green-fluorescent protein (GFP) (NCTC 12023)^52^ and *Escherichia coli* Nissle 1917^53^ (EcN) were grown overnight in LB (Sigma) and the O.D was measured on the following day at 650nm. Aliquots corresponding to 10^9^ particles of bacterial were centrifuged at 6000 rpm for 5 minutes and the pellet was washed twice in the fresh LB then twice in PBS (Sigma). Cells were then heat-killed at 65°C for 30 minutes and stored in aliquots at -20°C till further use. For monocyte stimulation assays, 0.1×10^6^ MACS-purified CD14^+^ monocytes (Mitenyi Biotec) were cultured in 150 μl medium in duplicates in round bottom 96-well plates and exposed to combinations of ST (monocyte to bacteria ratio = 1:2), EcN (monocyte to bacteria ratio = 1:2), ultrapure 200 ng/ml LPS (Enzo Life Sciences; Cat.# ALX-581-008), 10 μg/ml anti-IL-10R (Biolegend; clone: 3F9), 10 μg/ml anti-IL-10 (R&D; Cat.# MAB217; clone: 23738), 10 μg/ml anti-IL-1R1 (R&D; Cat.# AF269; polyclonal Goat IgG) for 16 hours (the last 4 hours of culture in the presence of BFA (Sigma) in the case of intracellular staining) in complete RPMI with L-glutamine (Sigma) supplemented with 10 % human serum (Sigma; Cat.H4522), non-essential amino acids (Gibco); 1 mM Sodium-Pyruvate (Gibco) and 100 U/ml penicillin and 10 μg/ml streptomycin (Sigma).

### Protein level analysis

Cytokine levels in the supernatants were assayed using the Milliplex human cytokine/chemokine magnetic bead 41-plex panel (Milliplex Billerica, MA, USA; HCYTOMAG-60K-PX41) and acquired on a Luminex LX200 flow reader. IL-23p19 cytokine level was assayed using a MSD (Meso Scale Discovery) in-house assay. In short, an IL-23p-19 selective antibody (Ab1, clone #32) was biotinylated following the Biotin, EZ-Link™ NHS-PEG4-Biotin instructions (ThemoScientific, Cat.# 2161299). For detection, an anti-p40 antibody (Ab-2, clone #42) was sulfotaged according to manufactures instructions (MSD Gold Sulfo-Tag NHS; Cat.# R91A0-1 and MSD Gold Sulfo-Tag NHS-Ester Conjugation Packs; Cat.# R31AA-1). MSD Streptavidin Gold plates (MSD; Cat.# L15SA-1) were washed, blocked, coated with biotinylated Ab-1, and washed according the manufacture instruction. 50 ul of supernatant or standard were added to the plate and incubated overnight at 4 °C while shaking. Plates were washed 3 times and detection antibody was added and incubated for 2 hrs while shaking at room temperature. Plates were washed and read buffer added. Plates were read with a MSD reader Quick Plex S120.

### Surface marker analyses by flow cytometry

The expression of cell surface markers was analysed by staining for 15 min at room temperature (RT) in PBS supplemented with 0.5% (v/v) human serum (FACS buffer). To allow exclusion of dead cells, cells were stained prior to fixation using Fixable Viability Dye eFluor® 780 (eBioscience). Following fluorophore-conjugated antibodies were used for analysis (supplier, clone): anti-CD3 (BD Biosciences; UCHT1), anti-CD4 (Biolegend; RPA-T4), anti-CD8 (Biolegend; SK1), anti-CD14 (Biolegend; M5E2), anti-CD16 (Biolegend; 3G8), anti-CD19 (BD Biosciences; SJ25C1), anti-CD25 (Biolegend; MA-A251), anti-CD45RA (Biolegend; HI100), anti-CD56 (BD Biosciences; NCAM16.2), anti-HLA-DR (Biolegend; L243). For combined analysis of surface marker and intracellular cytokines, surface-stained cells where subsequently stained as described in the section Intracellular cytokine staining. Data were acquired on a LSRII flow cytometer (BD Biosciences).

### Intracellular cytokine staining (ICCS)

Cells were stimulated in the presence of brefeldin A (BFA, 10 µg/ml, all from Sigma) for the final 4 hours of cell culture. Cells were fixed and permeabilised with Cytofix/Cytoperm (BD Biosciences) according to the manufacturer’s instructions. To exclude dead cells from the analysis, cells were stained prior to fixation using Fixable Viability Dye eFluor® 780 (eBioscience). Following fluorophore-conjugated anti-cytokine antibodies were used for analysis (supplier; clone): anti-IL-1α (Biolegend; 364-3B3-14), anti-IL-1β (Biolegend; H1b-98), anti-IL-4 (BD Biosciences; MP4-25D2), anti-IL-6 (eBioscience; MQ2-13A5), anti-IL-10 (Biolegend; JES3-9D7), anti-IL-13 (eBioscience: 85BRD), anti-IL-17A (eBioscience; eBio64DEC17), anti-IL-23p19 (eBioscience; 23dcdp), anti-GM-CSF (BD Biosciences; BVD2-21C11), anti-IFN-γ (eBioscience; 4S.B3), anti-TNF (Biolegend: MAb11). For the detection of IL-12p40 the combination of biotin-conjugated anti-IL-12p40 (BD Biosciences; C8.6) and streptavidin STV-CF594 (BD Biosciences; Cat.# 562318) were used. Data were acquired on a LSRII flow cytometer (BD Biosciences).

### Monocyte cluster validation and tSNE analysis of flow cytometry data

For the validation of scRNA-seq-identified monocyte clusters at the protein level, cells were stained for the following marker (supplier; clone): anti-CD14 (Biolegend; M5E2), anti-HLA-DR (Biolegend; L243), anti-CCL20/MIP-3 alpha (R&D; 67310), anti-CCL20/MIP-3 alpha (R&D; 114906), anti-IL-23p19 (eBioscience; 23dcdp), anti-CCL2 (eBioscience; 5D3-F7), anti-RPS6 (R&D; 522731), anti-SPINK1 (R&D; 839305), anti-IDO-1 (eBioscience; Eyedio), EBI3 (eBioscience; ebic6), anti-S100A8 (Invitrogen: CF-145), anti-S100A9 (Biolegend, MRP-14), anti-TNF (Biolegend: MAb11). Unconjugated antibodies targeting RPS6 and SPINK were conjugated in house using the Expedeon (Innova Biosciences) Lightning–Link PE/Texas Red or DyLight 405 antibody labelling kit according to the manufacturer’s instructions, respectively. t-Distributed Stochastic Neighbor Embedding (TSNE)-based analysis was executed on FCS files compensated for spillover between channels and gated on live CD3^−^ CD19^−^CD56^−^CD14^+^ single cells, down-sampled to 3.000 cells per sample. A single FCS file was generated by concatenating individual samples FCS files prior to tSNE unsupervised analysis using the FlowJo (Treestar) tSNE plugin54, 55 based on cluster defining genes. The following settings were used: Iterations = 1000; Perplexity = 200; Eta = 20; Theta = 0.5. and included the following parameters: CD14, HLA-DR, CCL2, CCL20, IL-23p19, IDO-1, RPS6, SPINK1, S100A8 and TNF. IL-23^+^ inflammatory monocytes were defined by combined IL-23p19, CCL20 and TNF expression, IFN-induced monocytes were defined by combined HLA-DR and IDO-1expression; microbicidal monocytes were defined by combined CCL2, S100A8 and CD14 expression and apoptotic monocytes were identified by combined RPS6 and SPINK-1expression).Each individual analysis was performed on samples that were stained and acquired (LSRII (BD Biosciences)) on the same day.

### Analysis of cytokine-induced STAT3 phosphorylation by flow cytometry

Whole blood (Δ*IL10RA*) or PBMC (Δ*IL10RB*) were surface stained with anti-CD3-PE-Cy5 or anti-CD3-FITC (Biolegend; clone: UCHT1), anti-CD4-BV605 (Biolegend; clone: OKT4), anti-CD8-PE-CF594 (BD Biosciences; clone: RPA-T8), anti-CD14-BV650 (Biolegend; clone: M5E2), and anti-CD19-BV711 (BD biosciences; clone: HIB19 or clone: SJ25C1) for 20 min. Five minutes through surface staining, cells were stimulated with 100 ng/ml human recombinant IL-10 or IL-6 (Peprotech) for 15 min. Cells were subsequently fixed in pre-warmed (37°C) BD Cytofix for 10 minutes (BD biosciences) at 37°C. After fixation, cells were permeabilised on ice with ice-cold Perm Buffer III (BD biosciences) and stained with anti-pSTAT3 (pY705)-Alexa Fluor 647 (4/P-STAT3, BD Phosflow) for 1 h at room temperature before acquiring on a LSRII flow cytometer (BD Biosciences). For the analysis of IL-10 responsiveness in stimulated monocytes, cells were washed (2 times) in RPMI (Sigma) and incubated in RPMI for 2 hours. Following incubation, cells were washed (2 times) in RPMI and stimulated for 15 minutes in complete RPMI with 50 ng/ml recombinant human IL-10 (Preprotech). Cell were then stained in PBS (Sigma) on ice with Fixable Viability Dye eFluor® 780 (eBioscience). Subsequently, cells were fixed, permeabilised and stained as described above.

### IL-10 and IL-23 ELISpot assay

ELISpot plates (Merck; Cat.#MSHAS4510) were incubated with sterile PBS (Gibco) and coated over night at 4°C with anti-IL-12/IL-23p40 (Mabtech; Cat.# MT86/221; 10 µg/ml in PBS) and anti-IL-10 (Mabtech; Cat.# 9D7;15 µg/ml in PBS) antibodies using 100 µl/well. On the next day plates were flicked and blocked using 100 µl/well 1x sterile PBS (Gibco) supplemented with 10% FCS (Sigma) for 2 hours at 37°C. Following, the buffer was removed and MACS-purified monocytes were plated at 2-fold concentration steps ranging from 6250 to 50.000 cells/well in complete RPMI medium supplemented with 10% FCS. Stimulation and incubation was performed over night at 37°C and 5% CO2 using combinations of LPS (200 ng/ml) and IL-10R blocking antibody (10 µg/ml). Following 16 hours incubation, cells were removed by flicking the plate and wells were washed extensively 10 times, each 1 minute incubation, with 200 μl washing buffer (0.1% Tween20 (Sigma) in PBS (Gibco). Secondary antibodies were diluted in PBS supplemented with 0.5% FCS (anti-IL-IL-23p19 (Mabtech; Cat.#/clone: MT155-HRP; 1 μg/ml) and anti-IL-10-biotin (Mabtech; Cat.#/clone: 12G8: 1 μg/ml). 100 μl/well antibody mixture was added to each well and incubation was performed for 2 hours at 37°C. Plates were then washed extensively 10 times, each 1 minute incubation. For the development of spots first 100 μl/well substrate solution (BCIP/NBT; Sigma) was added and development continued until distinct blue/grey spots emerged (approximately 5 minutes). Plates were again extensively washed 10 times, each 1 minute incubation, with 200 μl washing buffer. Then 100 μl/well substrate solution (AEC; BD Pharmingen) was added to each well and development was continued until distinct red spots emerged (approximately 5 minutes). Plates were left to dry in the dark and images were acquired using an AID EliSpot Reader Systems. The anti-IL-10 clone 12G8 was conjugated with biotin (Novus Biologicals Lightning-Link Rapid Type A Biotin Antibody Labeling Kit).

### Generation and pre-processing of single-cell RNA-sequencing data

CD14^+^ monocytes were purified from PBMC by MACS positive selection (Mitenyi Biotec) according to the manufacturers’ instructions from 2 human donors. The purity of sorting and viability was assessed by surface staining and flow cytometry (anti-CD14 (Biolegend; M5E2); Fixable Viability Dye eFluor® 780 (eBioscience). Cells were plated in 96-well U-bottom plates at a cell density of 0.5×10^6^ cells/well in 200 μl complete RPMI and were left unstimulated or exposed to combinations of ultrapure 200 ng/ml LPS (Enzo Life Sciences; Cat.# ALX-581-008) and 10 μg/ml anti-IL-10R (Biolegend; clone: 3F9) for 16 hours. For droplet-based single cell RNA-sequencing analysis cells were collected in RPMI with L-glutamine (Sigma) supplemented with 0.5% human serum (Sigma; Cat.# H4522) and the cell number was adjusted to 1000 cells/μl. Single cell suspensions were kept on ice, washed in PBS with 0.04% BSA and resuspended. 10.000 single cells/channel were captured in droplets on Chromium 10x Genomics platform (less than one hour following the termination of stimulation). Library generation for 10x Genomics v2 chemistry was performed following the Chromium Single Cell 3ʹ Reagents Kits User Guide (CG00052). Quantification of library was performed using an Agilent Bioanalyzer and Bioanalyzer High Sensitivity DNA Reagents (Cat.# 5067-4627). Single-cell RNA-sequencing libraries were generated using the 10x Genomics Single Cell 3’ Solution (version 2) kit and sequenced to a minimum mean depth of 44.3k reads/cell (Illumina HiSeq 4000). An average of 2850 cells/per sample and 1150 genes/cell were recovered. Data analysis was performed using Python3 pipelines (https://github.com/sansomlab/tenx) written using the CGAT-core library #https://doi.org/10.12688/f1000research.18674.1. Read mapping, quantitation and aggregation of sample count matrices was performed with the 10x Genomics Cell Ranger pipeline (version 2.1.1) and version 1.2.0 (GRCh38) reference sequences. No normalisation was applied during the aggregation step. Cells with barcodes common to samples sequenced on the same lane(s) were removed from the analysis to avoid issues associated with index hopping. Aggregated count matrices were randomly down-sampled in order to normalise the median number of UMIs per-cell between the samples (“downsampleMatrix” function from the DropletUtils R package). Down-sampling was performed separately for the within-donor cross-condition analysis (Figure 3) and the cross-donor analysis of LPS + anti-IL10R stimulated monocytes (Supplementary Fig. 4b).

### Cross-condition analysis of single-cell RNA-sequencing data from a single donor

The dataset was filtered to remove cells with <500 genes, >5% mitochondrial reads per cell or that expressed lymphocyte markers (*CD3*, *CD79A* or *CD79B*, n=33 cells) leaving a total of n=2420 unstimulated, n=3149 LPS stimulated and n=2273 LPS + anti-IL-10R stimulated monocytes. Per-cell UMI counts were then normalised, scaled and the effects of total UMI counts, percentage of mitochondrial counts and cell cycle (“all” effects; using known G2 and S phase associated genes^56^) regressed out with the Seurat R package (version 2.3.4).

Significantly variable genes (n=1164) were selected using the “trendVar” function from the R Bioconductor package Scran (minimum mean log-expression 0.05, BH adjusted p-value < 0.05). These genes were used as input for principal component analysis (PCA), and significant PCs (n=33) identified using Seurat (“JackStraw” test, BH adjusted p < 0.05). These principle components were used as input for alignment with Harmony (version 0.0.0.9, theta = 0). The first 30 Harmony components were used for graph-based clustering as implemented in Seurat (“original” Louvain algorithm, resolution=0.5). Significant cluster markers were identified using the Seurat “Findmarkers” function (Wilcoxon tests, BH adjusted p value < 0.05). The tSNE projection (Figure 3a) was computed using the 30 Harmony components and a perplexity of 50. Geneset over-representation analysis of cluster marker genes (Supplementary Fig. 4a) was performed using one-sided Fisher’s exact tests (as implemented in the “gsfisher” R package https://github.com/sansomlab/gsfisher) with Biological Process gene sets obtained from the Gene Ontology (GO) database^57^. For this analysis cluster-specific gene universes were defined as those genes expressed in 10% percent of cells (either within or outside the cluster of interest). Genesets were independently filtered for an odd ratio of ≥ 1.5 and n ≥ 3 foreground genes before multiple testing correction (enrichments with a BH adjusted p < 0.05 were considered significant).

### Comparison of LPS + anti-IL-10R single-cell RNA-sequencing data from two donors

The dataset was filtered as for the cross-condition analysis (removing n=23 contaminating lymphocytes) and an equal number (n=2280) of cells randomly sub-sampled from each donor for further analysis. Normalisation and scaling (including regression of cell-cycle effects), identification of variable genes (n=849), identification of significant PCs (n=35) and clustering (resolution=0.6) was performed as described above (no alignment performed).

### Identification and characterisation of gene co-expression modules in the RISK cohort

RPKM expression values for the RISK cohort^20, 21^ were retrieved from GEO (GSE57945), upper-quartile normalised and log2(n+1) transformed. Prior to WGCNA analysis the dataset was filtered to retain (n=13,850) genes that had a transformed expression value ≥ 1.5 in > 10% of the patients. The WGCNA R package (version 1.66) was then applied as follows (i) data was cleaned using the “goodSamplesGenes” function (with parameters: minFraction=0.5, minNsamples=4, minNGenes=4, minRelativeWeight=0.1), (ii) outlying samples (n=11) were identified and removed using the “cutreeStatic” function (with parameters: cutHeight=110, minSize=20), (iii) adjacencies calculated with the “adjacency” function (parameters: type=signed_hybrid, power=5, corFnc=cor, corOptions use=p and method=spearman”), (iv) the topological overall matrix computed with the “TOMsimilarity” function (parameter TOMType=unsigned), (v) a dynamic tree cut performed with function “cutreeDynamic” (parameter deepSplit=2, pamRespectsDendro=FALSE, minClusterSize=30) and (vi) modules merged with the “mergeCloseModules” function (parameter cutHeight=0.25). Genes assigned to modules were subject to geneset over-representation analysis as described for the single-cell RNA-sequencing analysis with gene categories obtained from the Gene Ontology (GO) database^58^, KEGG pathways^59^ and sets of cell type marker genes (xCell)^60^. Indicative monikers for the modules were manually assigned based on genes with high module membership (r > 0.8 with the module eigengene) as well as GO category, KEGG pathway and xCell marker enrichments (see Suplementary Table 4). The overlap of the identified WGCNA modules with IBD GWAS associated genes was performed using the associations reported by Ellinghaus *et. al*.^51^ (Ellinghaus Supplementary Table 3a, n=156 genes) or by de Lange *et. al*.^5^ (de Lange Supplementary Table 2). The associated genes from de Lange *et. al*. were conservatively filtered to retain only “implicated” genes or genes that originate from single-gene loci (n=89 genes, 48 of which overlap with the Ellinghaus *et. al*. set).

### Identification and characterisation of an IL-10-responsive monocyte gene signature

The IL-10 regulated inflammatory monocyte gene signature (Figure 5e) was constructed from (i) genes that showed ≥ 3-fold higher expression in at least one of the three LPS + aIL10R clusters than was found in any of the unstimulated or LPS-stimulated clusters (single-cell RNA-sequencing analysis, Figure 3a), (ii) genes that showed significantly higher expression in the CD14+ positive PBMC fraction upon with anti-IL-10R + LPS stimulation versus LPS stimulation alone (n=1, IL-1B, Supplementary Fig. 5), and (iii) two genes from the literature: CD14, a well-established monocyte type cell marker and OSM which is known to be expressed by inflammatory monocytes in human IBD and to drive colitis in the Hh + anti-IL-10R model^18^. The predictive ability of genes in the IL-10 regulated inflammatory monocyte signature was investigated in RNA-seq data from the RISK cohort and Affymetrix microarray data from the Janssen^22^ (UC-cohort GSE12251, Figure 5f). The UC-cohort GSE12251 includes data from patients with active UC that had a total Mayo score between 6-12 and an endoscopic sub-score of at least 2. The response to infliximab was defined as endoscopic and histologic healing in patients that underwent a second flex sigmoidoscopy with rectal biopsies 4 weeks after the first infliximab infusion in case of a single infusion and at 6 weeks if they received a loading dose of infliximab at weeks 0, 2 and 6. Microarray datasets from the Janssen (GSE12251) and Arijs (GSE16879)^23^ cohorts were RMA normalised (R Bioconductor package “oligo”) and annotated with data from the R Bioconductor “hgu133plus2.db” package (signals from probes mapping to the same gene symbol were averaged). Predictive ability was assessed by computation of area under the precision recall curve (AUPRC) using the R PRROC library.

### Real-time PCR

Real-time PCRs were performed in 96-well plates using the PrecisionPLUS qPCR Mater Mix (Primer Design) and the CFX96 Touch Real-Time PCR Detection System machine (BIO-RAD). The expression of transcripts was normalized to expression of Large Ribosomal Protein (RPLPO). Data analysis was performed using the Cycles threshold (ΔΔCt) method and expressed as mRNA relative expression ΔΔCt. The following TaqMan probes (Applied Biosystems) were used for qPCR analysis: IL23A (Hs00900828_g1), IFNG (Hs00989291_m1), RPLPO (Hs99999902_m1).

### Gene array expression and statistical analysis

Background correction and quantile normalization of the gene expression data obtained from the Affymetrix Human Transcriptome Array 2.0 (HTA2) platform were performed by the RMA method^16^. Each gene’s expression was defined at the level of the “transcript cluster”, which is the collection all probesets on the HTA2 platform that are located within a gene’s annotated position in the genome. These gene positions were defined by the Affymetrix NetAffxTM NA34 release, which is based on the GRCh37 human reference genome. Differential expression at the gene-level was performed using the empirical Bayes method implemented in LIMMA^61^ controlling for donor in the model. Differential expression between a given stimulation vs. PBMC control contrast was considered at a Benjamini-Hochberg adjusted p-value < 0.05 and absolute fold change > 2. A heatmap was produced using the heatmap.2 function from the gplots package (R-3.4.2) to visualise the union of genes that were regulated in each stimulation vs. PBMC control contrast.

### Mathematical modelling of cytokine interactions

We define a system of ordinary differential equations of the form 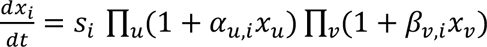, with *x_i_* the concentration of TNF, IL-1α, IL-1β, IL-6, IL-10 or IL-23 over time, *s_i_* the contribution of LPS, and α*_u,i_*(*β_u,i_*) the positive (negative) effect of cytokine *u* on the production rate of cytokine *i*. With additional time and/or dose dependent data, the model could be extended by including saturation of cytokine production (e.g. using Michaelis-Menten or Hill functions) or cytokine degradation. By minimizing the log likelihood, we fit the right-hand side of the equations to 1750 experimental data points, representing cytokine production rates by monocytes 16 hours after LPS stimulation. These rates were determined by intracellular flow cytometry in PBMC derived from healthy individuals, in the presence or absence of anti-IL-10R, and TNF, IL-1β, IL-6, IL-10, IL-23, IFN*γ*, anti-IFN*γ*, anti-IFNR1, anti-IL-1α, anti-IL-1β, anti-IL-1R1, anti-IL-6R, or anti-TNF (n=3-17). The frequency of cytokine (i.e. TNF, IL-1α, IL-1β, IL-6, IL-10 or IL-23) producing live CD14^+^ cells multiplied by the mean fluorescence intensity of the respective cytokine producing cell population was taken as a measure of cytokine production rate. All possible network configurations using a set of edges representing significant (paired Wilcoxon test, p≤0.05) direct cytokine interactions derived from this dataset were considered, and for each number of edges, an optimal model configuration was found. We did not model all 2^31^-1 possible configurations explicitly, as the vast majority of configurations could be logically excluded based on a small subset of network configuration fits. It was found that all optimal models were nested, i.e. the list of optimal models can be obtained by iteratively adding edges to the model. As additional edges will always improve the model fit, we use the Akaike Information Criterion (AIC) to identify the fourteen-edge model as the model with an optimal balance between edge number and model fit. For two nested models that differ one edge, the AIC is equivalent to a likelihood ratio test (LRT, alpha=0.16) with one degree of freedom. Fitting and parameter identifiability analysis of the fourteen-edge model was done in the Matlab based modelling environment ‘Data2Dynamics’, using a deterministic trust-region approach combined with a multi-start strategy; data2dynamics.org^62^.

### Resources for data visualization

Gene and protein networks were visualized using Cytoscape 3.6.1: http://www.cytoscape.org/63. Heatmaps for gene expression data visualisation were generated using the GENE-E or Morpheus tool from the Broadinstitute: http://www.broadinstitute.org/cancer/software/GENE-E/; https://software.broadinstitute.org/morpheus/.

---

**Supplementary Figure 1:**
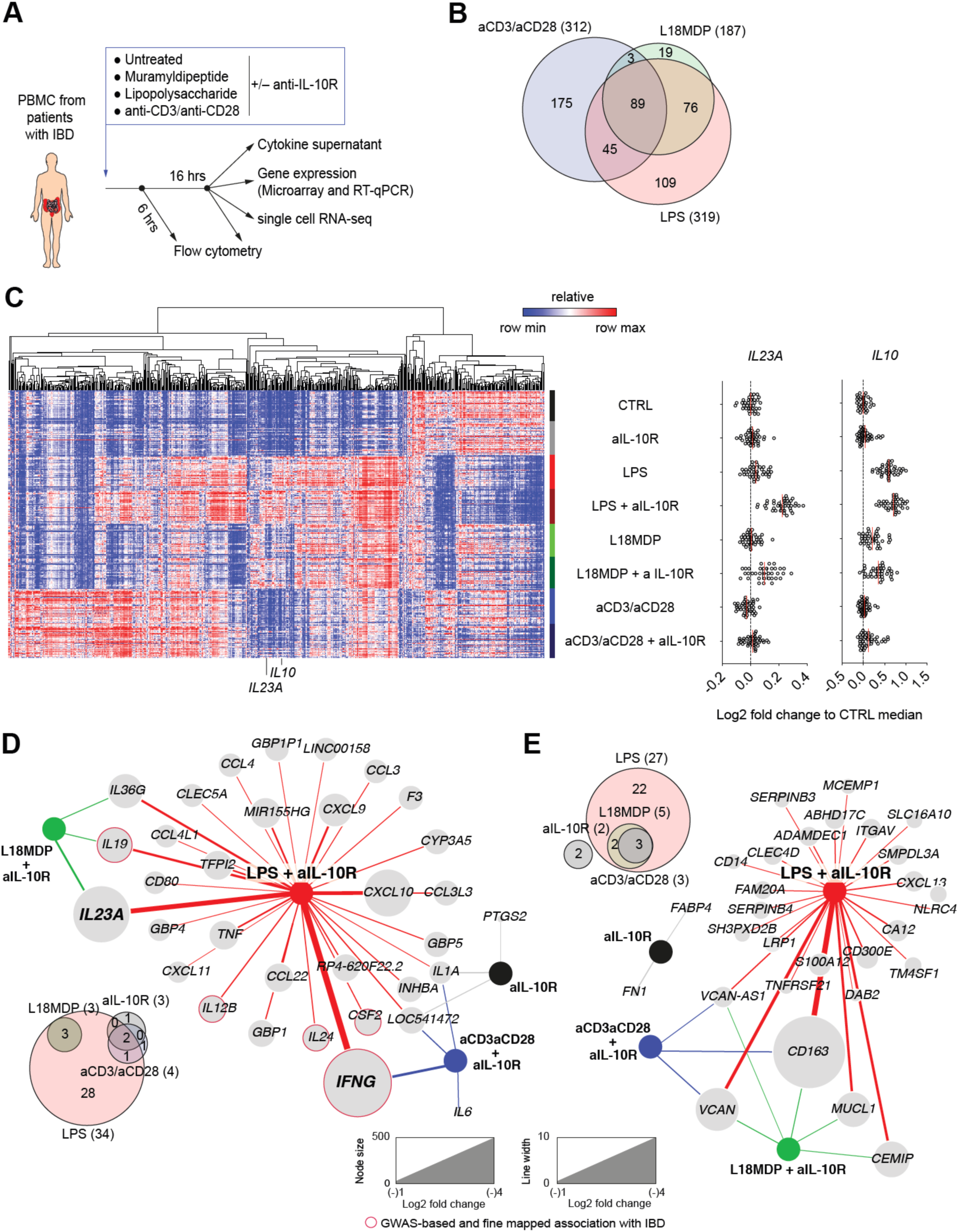
(**A**) Graphical illustration of *ex vivo* PBMC stimulations and experimental conditions. (**B**) Venn-diagram of differentially regulated genes in the three stimulation conditions (LPS, L18MDP, and αCD3/αCD28 coated beads). (**C**) Microarray analysis of patients with IBD (n=41; Supplementary Table 1) PBMC samples under diverse stimulation conditions. The heatmap shows transcripts found to be significantly differentially expressed (columns, fold change > 1.5, BH-adjusted p-value < 0.05) in the patient samples (rows) across diverse stimulation conditions (LPS, L18MDP and αCD3/αCD28) at 16 hours post stimulation. *IL23A* and *IL10* expression are shown on the right as log2 fold change expression to the unstimulated PBMC (CTRL) median expression. (**D**, **E**) Networks of genes significantly up-regulated (**D**) or downregulated (**E**) by IL-10 signalling blockade (aIL-10R) in unstimulated (black), LPS stimulated (red), L18MDP stimulated (green) and αCD3/αCD28 stimulated (blue) patient samples (fold change > 1.5; node size and edge-thickness are proportional to fold change).

**Supplementary Figure 2:**
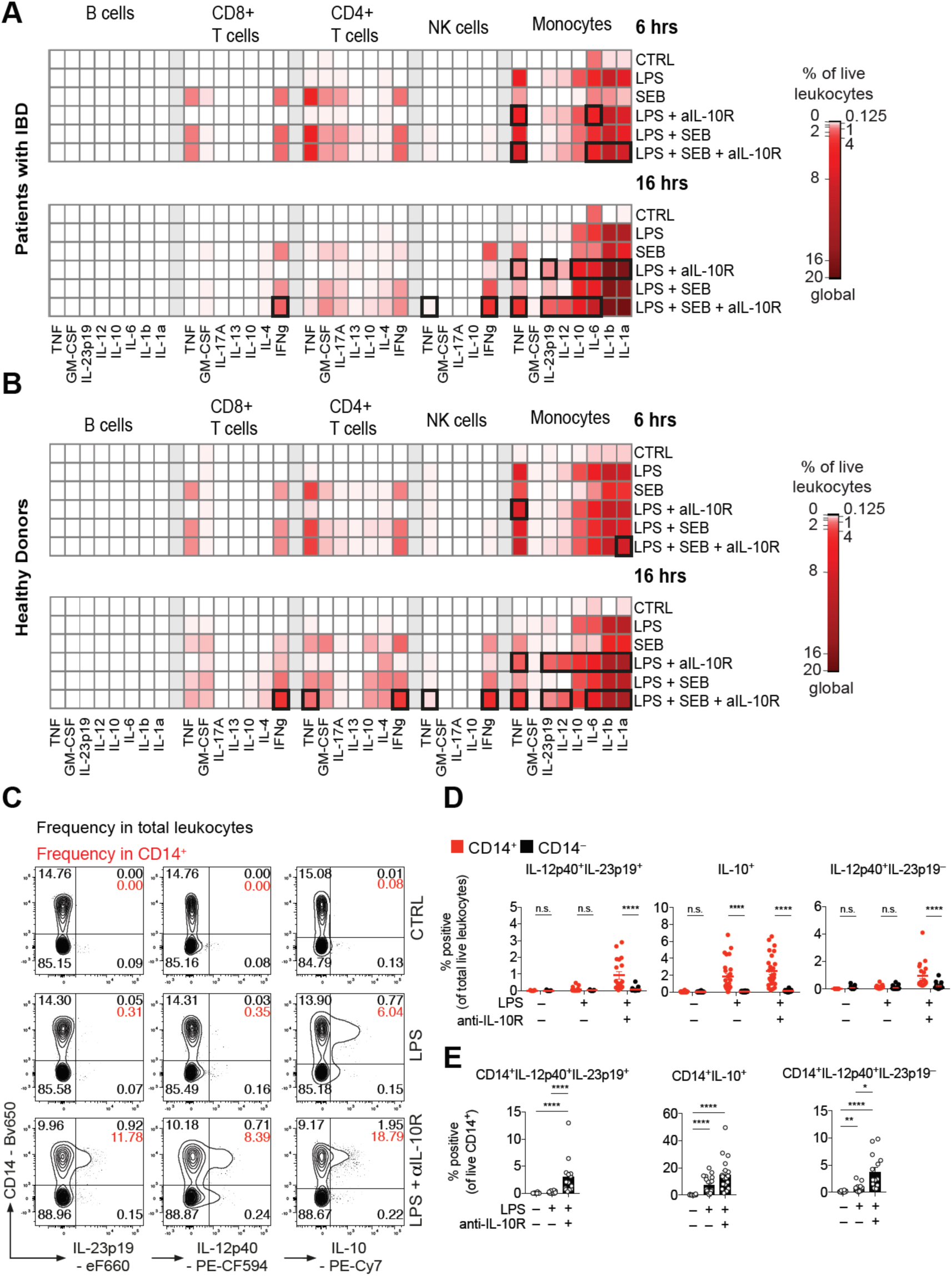
(**A** and **B**) PBMC from patients with IBD (A; n = 7) or health individuals (B; n = 10) were stimulated for 6 or 16 hours with combinations of LPS (200 ng/ml), SEB (1 μg/ml) and IL-10R blocking antibodies (10 μg/ml) with BFA present for the last 4 hrs of culture. Cells were then surface stained to identify distinct leukocyte populations followed by intracellular cytokine staining. The frequency of cytokine producing cells for each population was assessed and presented as % cytokine producing cells of total live leukocytes. Monocytes were identified as CD14^+^CD3^−^CD19^−^CD56^−^, CD4^+^ T cells as CD3^+^CD4^+^CD8^−^ CD14^−^CD19^−^CD56^−^, CD8^+^ T cells as CD3^+^CD8^+^CD4^−^CD14^−^CD19^−^CD56^−^ NK cells as CD56^+^CD3^−^CD14^−^CD19^−^ and B cells were identified as CD19^+^CD3^−^CD14^−^CD56^−^ (Mean; bold squares indicate p<0.05; Mann-Whitney test). (**C**) Contour plot presentation of IL-23p19-, IL-12p40- and IL-10-producing live leukocytes and CD14 surface expression measured at 16 hours post stimulation in HD PBMC. (**D**) Summary of frequencies of IL-12p40^+^IL-23p19^+^, IL-10^+^ and IL-12p40^+^IL-23p19^−^ CD14^+^ and CD14^−^ cells in HD total live leukocytes (n = 26). Mean +/– SEM; Mann-Whitney test. (**E**) Summary of frequencies of IL-12p40^+^IL-23p19^+^, IL-10^+^ and IL-12p40^+^IL-23p19^−^ of HD CD14^+^ cells (n = 26). Mean +/– SEM; Mann-Whitney test.

**Supplementary Figure 3:**
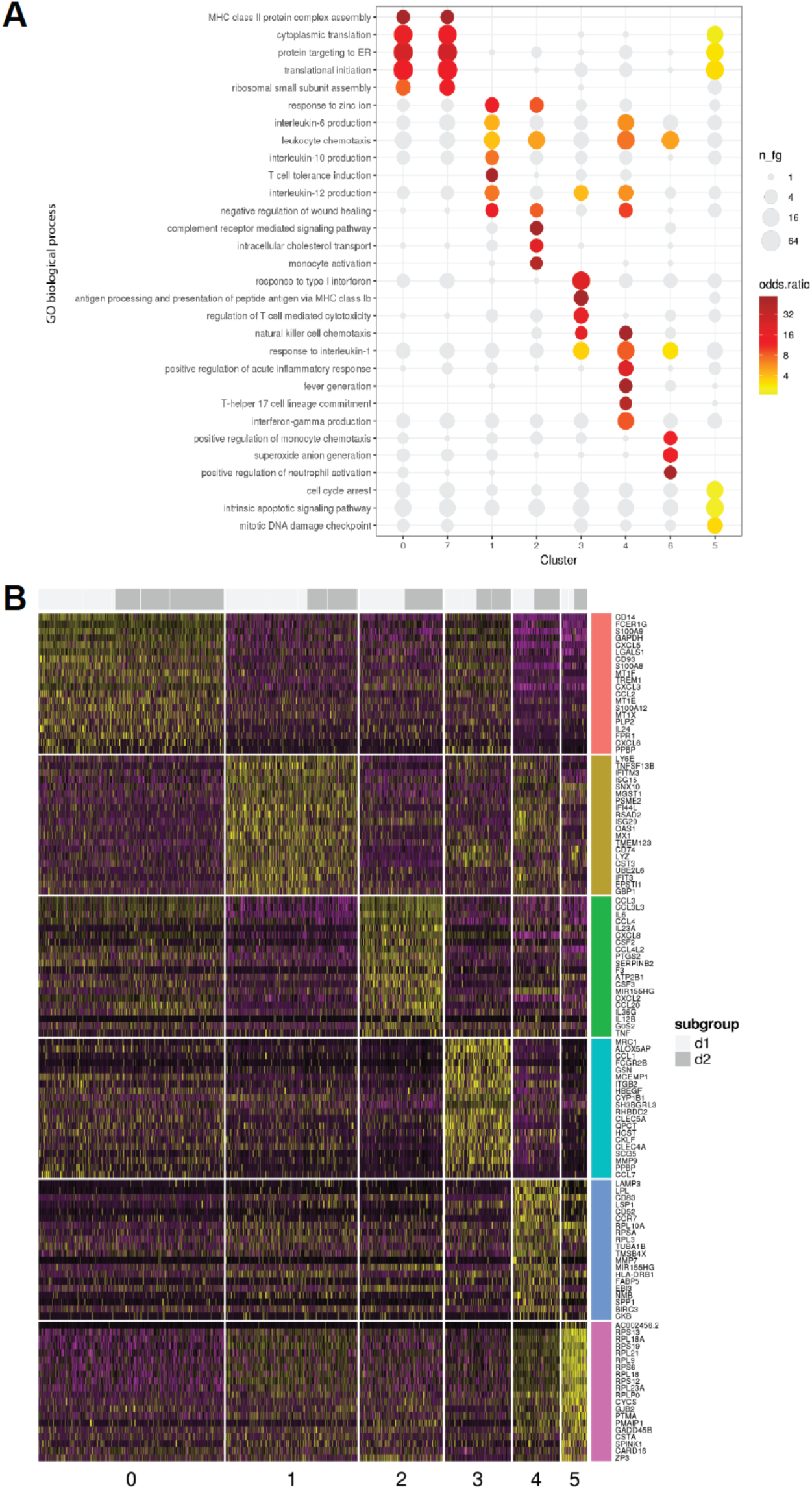
Single cell genomics identifies transcriptionally distinct monocyte clusters following combined LPS + anti-IL-10R stimulation. (**A**) Selected over-represented GO biological process gene sets of the 8 identified monocyte clusters across stimulation conditions (Unstimulated, LPS-stimulated and LPS + anti-IL-10R-stimulated). (**B**) Heatmap of cluster marker genes for cell-cycle corrected analysis of the LPS + anti-IL-10R condition alone (donors 1 and 2). Cluster 1 corresponds to cluster 3 in the aligned analysis (Interferon-induced monocytes), cluster 2 to cluster 4 in the aligned analysis (IL-23^+^ inflammatory monocytes) and cluster 3 to cluster 6 in the aligned analysis (MRC1^+^ microbicidial monocyte subpopulations).

**Supplementary Figure 4:**
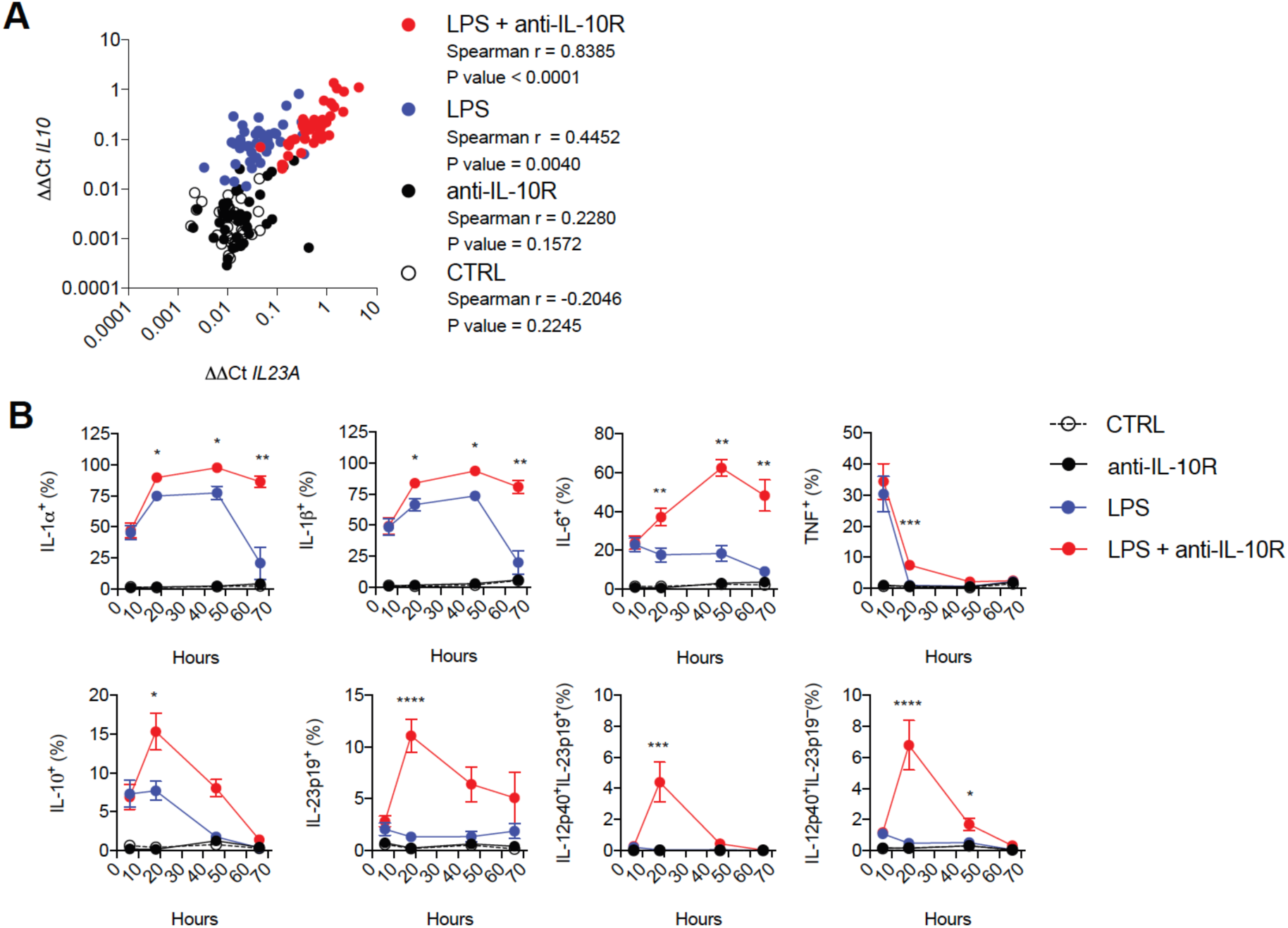
*IL23A* and *IL10* gene expression positively correlate under settings of LPS stimulation and IL-10 signalling blockade and monocytes produce cytokines with distinct kinetics. (**A**) RT-qPCR based ΔΔCt correlation analysis of *IL10* and *IL23A* gene expression in unstimulated, anti-IL-10R (10 μg/ml), LPS (200 ng/ml), and combined LPS and anti-IL-10R-stimulated PBMC (n=40). (**B**) PBMC were stimulated for up to 66 hours with combinations of LPS and anti-IL-10R. The frequencies of cytokine producing live CD14^+^ cells were assessed by surface and intracellular staining at different time points, as indicated (n=4-10, Mean +/– SEM; Kruskal-Wallis test; BH-adjusted p-values).

**Supplementary Figure 5:**
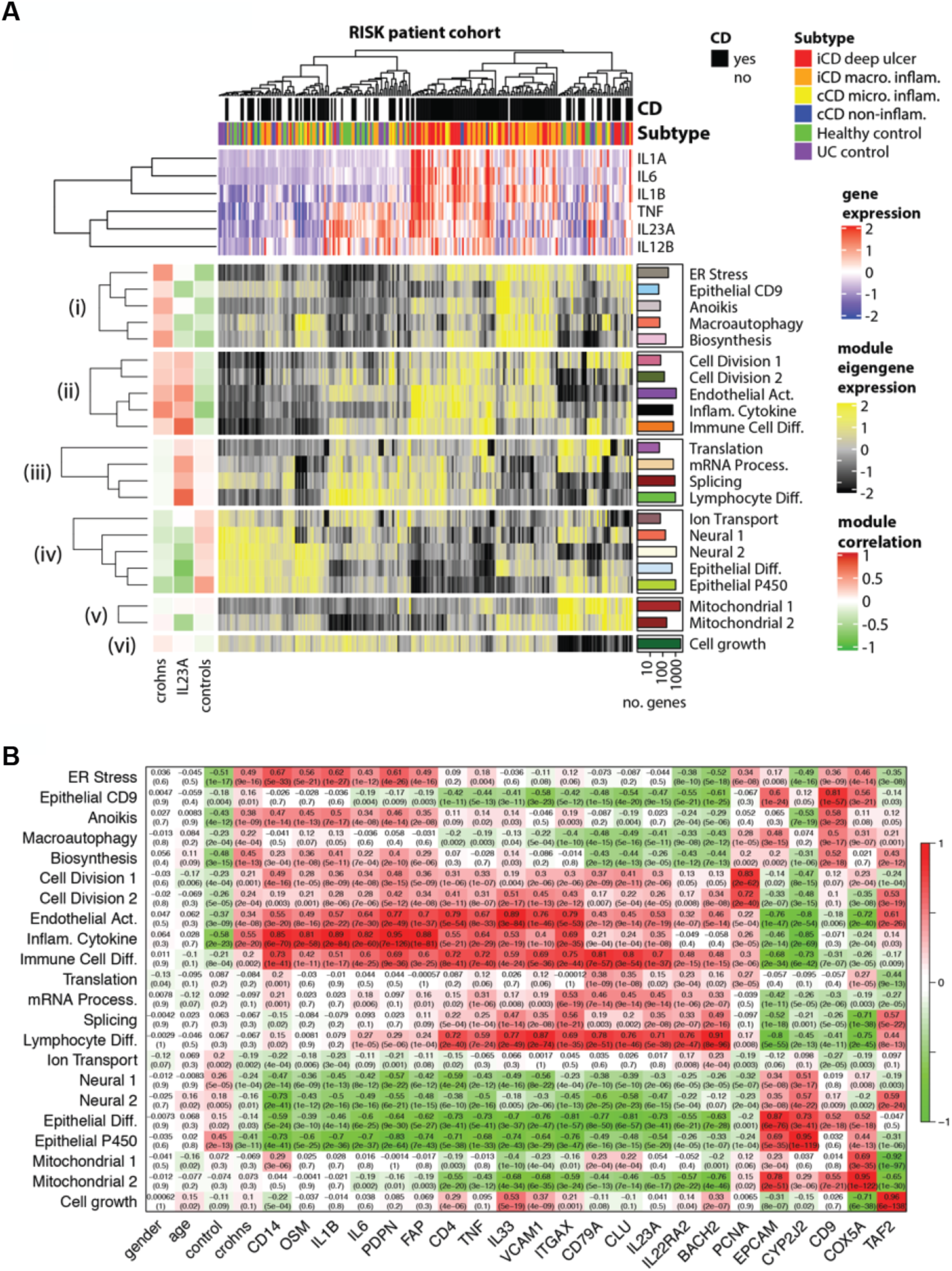
Identification of modules of co-expressed genes in patient biopsies from the paediatric RISK cohort. (**A**) The main heatmap shows expression of the eigen-genes of the n=22 identified modules of genes. Patients (columns) are hierarchically clustered by expression of the module eigen-genes. The heat map is annotated with (i) patient diagnosis and disease subtype (top panels), (ii) expression of key cytokines (upper heatmap), (iii) correlation of the eigen-genes with diagnosis of Crohn’s disease, *IL23A* expression and control status (left panel) and (iv) the numbers of genes within each of the modules (right panel). Modules were named according to enrichments for gene ontology and cell type genesets (Supplementary Table 4). (**B**) The heatmap shows the correlation of the RISK WGCNA modules (y-axis) with genes and traits of interest (x-axis).

**Supplementary Figure 6:**
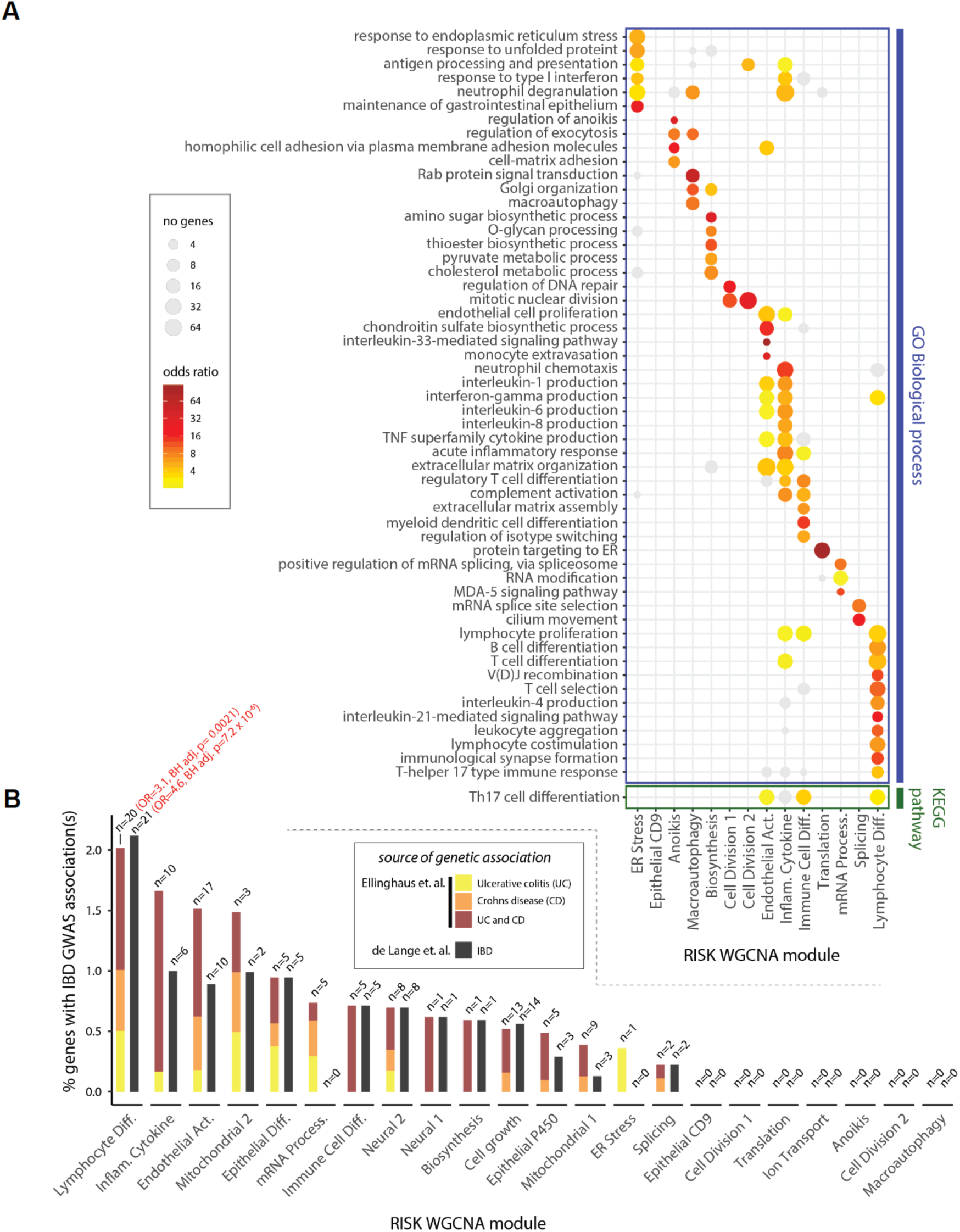
Characterisation of the RISK WGCNA modules. (**A**) The dot-plot shows the enrichment of selected biological process gene ontology categories and KEGG pathways in the WGCNA modules that were correlated with diagnosis of Crohn’s disease and/or *IL23A* expression (one-sided Fisher tests, BH-adjusted p<0.05, p-values adjusted separately for GO categories and KEGG pathways). (**B**) The frequency (y axis) of IBD GWAS-associated genes in the different RISK WGCNA modules (x axis). Genetically associated genes were sourced from Ellinghaus *et. al.*^51^ or de Lange *et. al.*^5^ (see methods). BH-adjusted p-values and odds ratios are given for significant enrichments (red text).

**Supplementary Figure 7:**
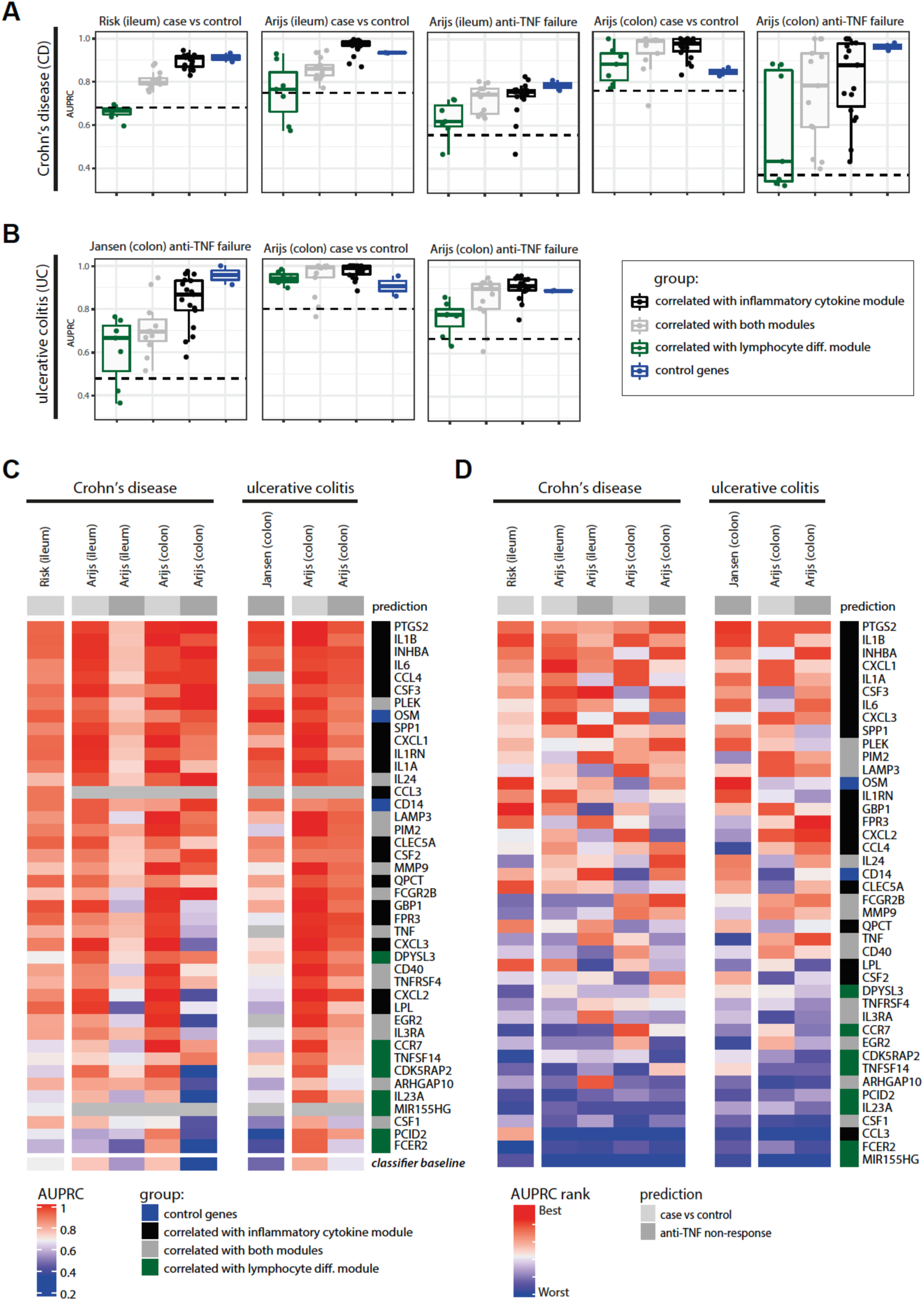
Ability of inflammatory monocyte signature genes to predict inflammatory bowel disease and non-response to anti-TNF therapy in three patient cohorts. Comparison of the ability of the identified subsets of monocyte genes (see Figure 4) to predict diagnosis of inflammatory bowel disease and TNF non-response in the RISK (GSE57945), Janssen (GSE12251) and Arijs (GSE16879) cohorts. (**A**) Predictions in Crohn’s disease with data from the RISK and Arijs cohorts. (**B**) Predictions in ulcerative colitis with data from the Janssen and Arijs cohorts. The panels from the RISK and Janssen cohorts in (A) and (B) are reproduced from Figure 4. The dashed lines indicate random classifier performance. (**C**) The heatmap shows the AUPRC values of the individual genes for the predictions shown in (A) and (B). (**D**) The heatmap show the ranks of the predictions for the individual genes. Genes in panels (C) and (D) are ordered by the mean of the AUPRC and AUPRC rank respectively. AUPRC: area under precision recall curve.

**Supplementary Figure 8:**
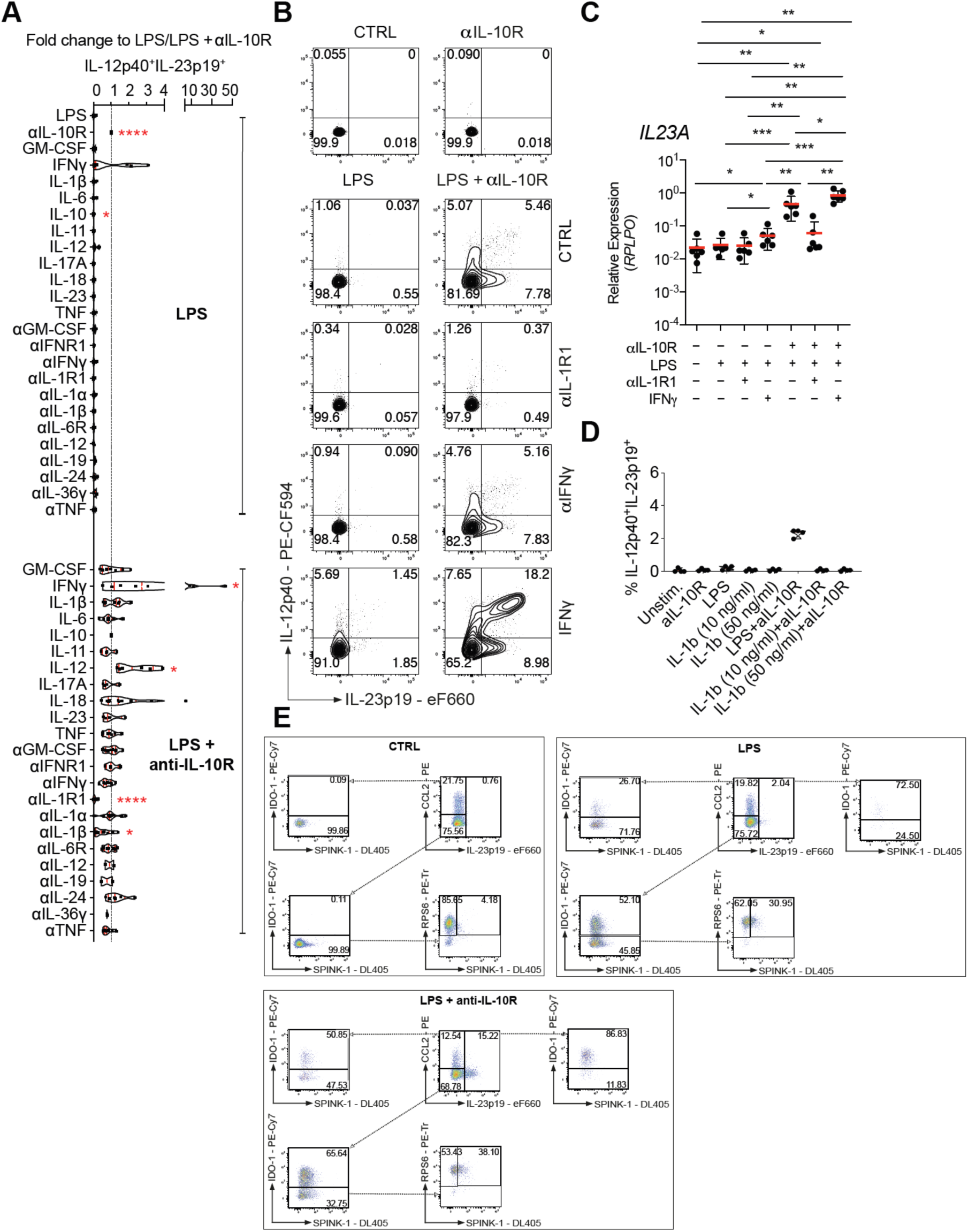
IL-1α and IL-1β signalling are essential for inducing monocyte IL-23 production. PBMC from patients with IBD (n > 4) were stimulated for 16 hours with combinations of LPS and αIL-10R in the presence of indicated exogenous human recombinant cytokine, cytokine receptor blockade or cytokine blockade. (**A**) The frequencies of cytokine producing live CD14^+^ cells were assessed by intracellular staining. Summary of IL-12p40^+^IL-23p19^+^ cell frequencies measured under diverse stimulation conditions; Wilcoxon test; 95% CI). (**B**) Representative dot plot showing intracellular IL-12p40 and IL-23p19 in CD14^+^ monocytes of one healthy donor. (**C**) RT-qPCR analysis of healthy donor PBMC stimulated with combinations of LPS, anti-IL-10R, anti-IL-1R1 and IFN-γ for 16 hours. (**D**) Frequencies of IL-12p40^+^IL-23p19^+^ live CD14^+^ monocytes after 16 hours stimulation with combinations of LPS (200 ng/ml), anti-IL-10R (10 μg/ml) and IL-1β (10 and 50 ng/ml). (**E**) Dot plot presentation of one representative experiment showing live CD14^+^-gated monocytes and sub-gating strategies based on the expression of IL-23p19, CCL2, IDO-1 RPS6 and SPINK-1 in non-stimulated, LPS-stimulated and combined LPS and anti-IL-10R stimulated PBMC.

**Supplementary Figure 9:**
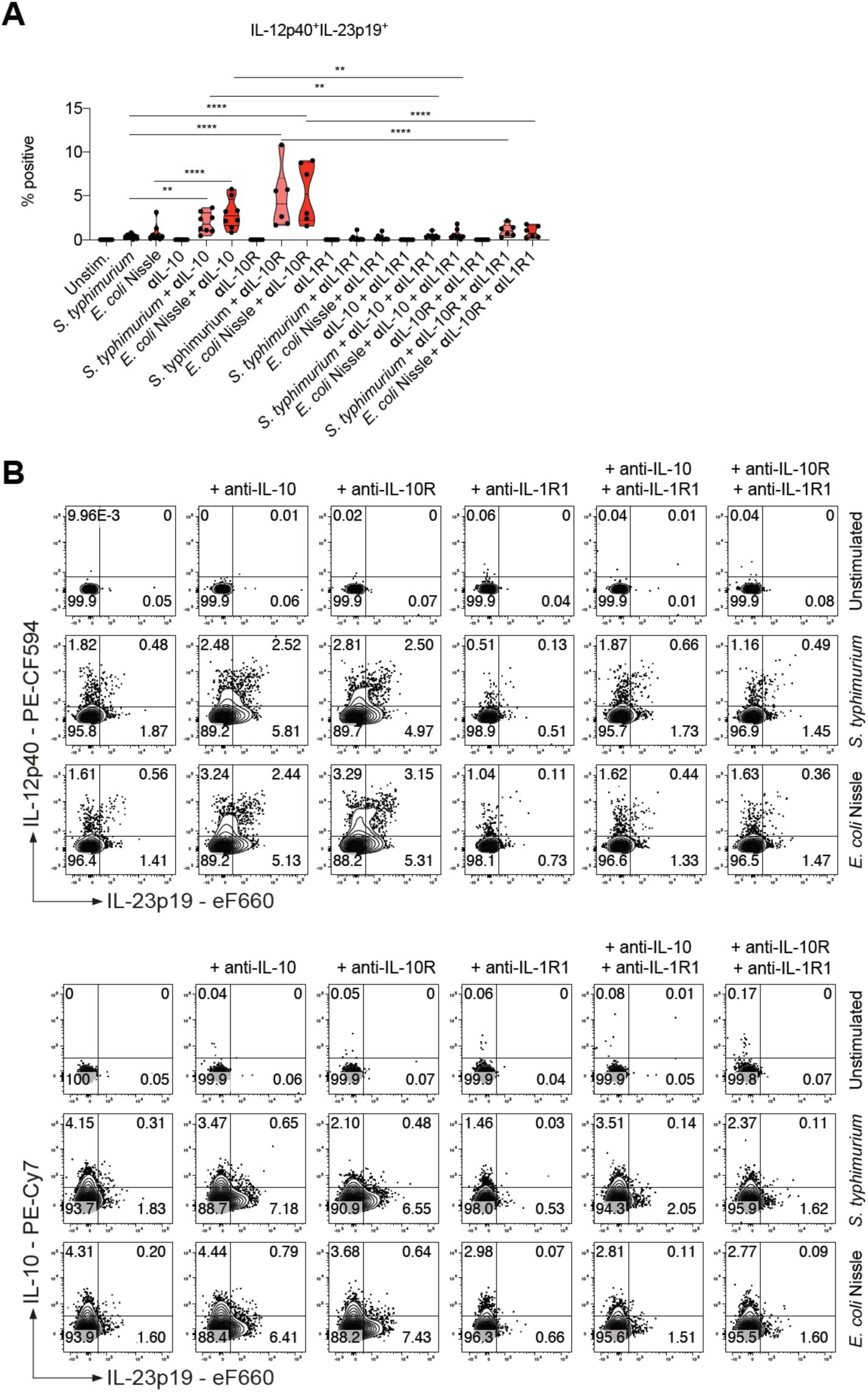
Monocyte IL-23 production induced by uptake of whole bacteria is dependent on IL-1R1 signalling. (**A**) Summary of frequencies of IL-12p40^+^IL-23p19^+^ live CD14^+^ cells as assessed by intracellular staining measured in indicated conditions at 16 hours following stimulation (n = 8, Mean +/– SEM; Kruskal-Wallis test; FDR-adjusted BH-adjusted p-values). (**B**) Representative dot plots showing intracellular IL-12p40, IL-23p19 and IL-10 in live CD14^+^ monocytes of one healthy donor.

## Notes

#### Summary of Updates

This version has been updated to increase the focus on the regulation of monocyte IL-23 responses in inflammatory bowel disease. We have added the analysis of a validation dataset (Supplementary Figure 7) to support our findings in the RISK patient cohort (Figure 4; Supplementary Figure 5 and 6).

