## Supplemental Table 1 for "Systems-level analysis of monocyte responses in inflammatory bowel disease identifies IL-10 and IL-1 cytokine networks that regulate IL-23"

Suppl. Table 1: IBD patients' characteristics

| | CD | UC | IBDu |
|---|---|---|---|
| Number | 27 | 12 | 2 |
| Age (Median, years) | 42 | 45 | 45 |
| Gender (M/F) | 13 M-14 F | 5 M-7 F | 1 M-1 F |
| Age of diagnosis (median; years) | 19 | 22 | 24 |
| (median, IQR) | | | |
| Disease Extent (%) | | | |
| E1 Proctitis | - | 1 | - |
| E2 Left sided | - | 4 | - |
| E3 Extensive | - | 3 | - |
| PSC | - | - | - |
| Crohn's disease | | | |
| Ileocolitis | | | |
| Ileitis | | | |
| Crohn's colitis | | | |
| Anti-TNFa (%) | 13% | - | - |
